# Bullying Victimization and Brain Development: A Longitudinal Structural Magnetic Resonance Imaging Study from Adolescence to Early Adulthood

**DOI:** 10.1101/2024.09.11.611600

**Authors:** Michael Connaughton, Orla Mitchell, Emer Cullen, Michael O’Connor, Tobias Banaschewski, Gareth J. Barker, Arun L.W. Bokde, Rüdiger Brühl, Sylvane Desrivières, Herta Flor, Hugh Garavan, Penny Gowland, Antoine Grigis, Andreas Heinz, Herve Lemaitre, Jean-Luc Martinot, Marie-Laure Paillère Martinot, Eric Artiges, Frauke Nees, Dimitri Papadopoulos Orfanos, Luise Poustka, Michael N. Smolka, Sarah Hohmann, Nathalie Holz, Nilakshi Vaidya, Henrik Walter, Gunter Schumann, Robert Whelan, Darren Roddy

**Affiliations:** Department of Psychiatry, Royal College of Surgeons in Ireland, Dublin 2, Ireland; Department of Child and Adolescent Psychiatry and Psychotherapy, Central Institute of Mental Health, Medical Faculty Mannheim, Heidelberg University, Square J5, 68159 Mannheim, Germany; German Center for Mental Health (DZPG), partner site Mannheim-Heidelberg-Ulm; Department of Neuroimaging, Institute of Psychiatry, Psychology & Neuroscience, King’s College London, United Kingdom; Discipline of Psychiatry, School of Medicine and Trinity College Institute of Neuroscience, Trinity College Dublin, Dublin, Ireland; Physikalisch-Technische Bundesanstalt (PTB), Braunschweig and Berlin, Germany; Social, Genetic and Developmental Psychiatry Centre, Institute of Psychiatry, Psychology & Neuroscience, King’s College London, United Kingdom; Institute of Cognitive and Clinical Neuroscience, Central Institute of Mental Health, Medical Faculty Mannheim, Heidelberg University, Square J5, Mannheim, Germany; Department of Psychology, School of Social Sciences, University of Mannheim, 68131 Mannheim, Germany; Departments of Psychiatry and Psychology, University of Vermont, 05405 Burlington, Vermont, USA; Sir Peter Mansfield Imaging Centre School of Physics and Astronomy, University of Nottingham, University Park, Nottingham, United Kingdom; NeuroSpin, CEA, Université Paris-Saclay, F-91191 Gif-sur-Yvette, France; Department of Psychiatry and Psychotherapy CCM, Charité – Universitätsmedizin Berlin, corporate member of Freie Universität Berlin, Humboldt-Universität zu Berlin, and Berlin Institute of Health, Berlin, Germany; German Center for Mental Health (DZPG), Berlin-Potsdam, Germany; Institut des Maladies Neurodégénératives, UMR 5293, CNRS, CEA, Université de Bordeaux, 33076 Bordeaux, France; Institut National de la Santé et de la Recherce Médicale, INSERM U A10 “Trajectoires développementales & psychiatrie”, University Paris-Saclay, Ecole Normale Supérieure Paris-Saclay, CNRS; Centre Borelli, Gif-sur-Yvette, France; AP-HP. Sorbonne Université, Department of Child and Adolescent Psychiatry, Pitié-Salpêtrière Hospital, Paris, France; Psychiatry Department, EPS Barthélémy Durand, Etampes, France; Institute of Medical Psychology and Medical Sociology, University Medical Center Schleswig Holstein, Kiel University, Kiel, Germany; Department of Child and Adolescent Psychiatry, Center for Psychosocial Medicine, University Hospital Heidelberg, Heidelberg, Germany; Department of Psychiatry and Psychotherapy, Technische Universität Dresden, Dresden, Germany; Centre for Population Neuroscience and Stratified Medicine (PONS), Department of Psychiatry and Psychotherapy, Charité Universitätsmedizin Berlin, Germany; Centre for Population Neuroscience and Precision Medicine (PONS), Institute for Science and Technology of Brain-inspired Intelligence (ISTBI), Fudan University, Shanghai, China; School of Psychology and Global Brain Health Institute, Trinity College Dublin, Ireland

**Author notes:** **<u>Corresponding author:</u>** Dr Darren Roddy, Senior Clinical Lecturer in Psychiatry, Royal College of Surgeons, Dublin 2.

## Abstract

This study investigated associations between bullying victimization and brain development using longitudinal structural MRI data from the IMAGEN cohort (n = 2,094; 1,009 females) across three time points (∼14, ∼19, and ∼22 years). A data-driven analysis revealed that higher bullying victimization was significantly associated with accelerated volumetric growth in subcortical and limbic regions, including the putamen (β = 0.12, 95% CI: 0.10–0.15), amygdala (β = 0.07, 95% CI: 0.05–0.09), hippocampus (β = 0.06, 95% CI: 0.04– 0.08), and anterior cingulate cortex (caudal: β = 0.05, 95% CI: 0.03–0.07; rostral: β = 0.06, 95% CI: 0.04–0.08). In contrast, bullying victimization was also significantly associated with reduced volumetric growth in the cerebellum (β = –0.09, 95% CI: –0.11 to –0.07), entorhinal cortex (β = –0.10, 95% CI: –0.13 to –0.07), and insula (β = –0.08, 95% CI: –0.11 to –0.06).Exploratory analyses indicated that females exhibited more pronounced changes in emotional processing regions, while males showed greater changes in motor and sensory areas. Overall, the findings indicate that bullying victimization is associated with widespread structural differences in brain development from adolescence to early adulthood, with sex-specific trajectories.

## Introduction

Bullying victimization, characterized by targeted and often chronic peer aggression— whether physical, verbal, or relational—is a common experience during childhood and adolescence, predominantly perpetrated by school peers (1). Persistent exposure to bullying is strongly associated with increased risk for depression, anxiety, suicidality, and impaired cognitive and social functioning (2-4). These adverse outcomes often extend into adulthood, raising concerns that bullying may be associated with disruptions in core neurodevelopmental processes during a critical period of brain maturation (5).

Adolescence is a particularly sensitive period for brain development, marked by extensive biological and psychological transformations (6). This phase involves substantial reorganization of neural circuits supporting executive function, emotion regulation, social cognition, and stress responsivity (7, 8). These changes are especially pronounced in frontolimbic regions (e.g., prefrontal cortex, anterior cingulate cortex), parietotemporal areas (e.g., superior temporal sulcus, temporoparietal junction), and subcortical structures (e.g., amygdala, hippocampus, striatum) (7, 8). While this plasticity supports adaptive development, it may also render the adolescent brain particularly vulnerable to adverse environmental exposures such as bullying victimization (5). Yet, how bullying victimization is associated with changes in brain development over time remains poorly understood.

Neurobiologically, experiences of bullying victimization may influence brain development through multiple interacting pathways (5). It has been linked to stress-induced neuroendocrine reactivity, neuromodulation, and limbic system dysregulation, which may collectively drive widespread structural brain alterations (5). These effects are mediated by multiple interconnected systems, including dysregulation of the hypothalamic-pituitary-adrenal (HPA) axis, heightened inflammatory responses, altered dopaminergic and serotonergic signaling, oxytocin pathway disruption, and imbalances in autonomic nervous system activity (9-13). Together, these systems can lead to sustained cortisol release and other neurochemical changes that result in downstream alterations in neurodevelopment, particularly in circuits involved in emotional, social, and cognitive functioning (5).

While cross-sectional structural MRI studies have offered evidence that bullying-related stress may alter brain structure (14), our understanding of its association with brain development over time remains limited. To date, only two longitudinal structural MRI studies have begun to address this question. Menken et al. (2023), using data from the ABCD study, found that children aged 9–11 exposed to bullying exhibited developmental differences,including steeper increases in hippocampal and entorhinal cortex volumes, along with accelerated cortical thinning in several frontal and temporal regions (15). Quinlan et al. (2020), using the IMAGEN cohort, reported that adolescents exposed to peer victimization from ages 14 to 19 exhibited steeper declines in left putamen volume, which predicted higher anxiety symptoms in early adulthood (16). However, their analysis focused on nine bilateral regions of interest, potentially overlooking broader patterns of neurodevelopmental change. While both studies were pioneering in establishing early longitudinal links between bullying victimization and brain development, each was limited to two imaging time points, restricting the ability to capture non-linear, region-specific developmental trajectories.

Indeed, brain development across adolescence is rarely linear (7, 8, 17). Cortical and subcortical structures often follow quadratic growth patterns, including periods of acceleration, deceleration, and stabilization (7, 8, 17). Two-time-point designs constrain our ability to capture such inflection points and may obscure meaningful developmental variability. In contrast, three-time-point longitudinal designs allow for more accurate mapping of non-linear, region-specific developmental trajectories, which is particularly important as the field moves toward the development of normative brain growth charts (18). Such charts may help identify neurodevelopmental shifts that contribute to diverging mental health outcomes, particularly in adolescents exposed to environmental stressors like bullying victimization.

To address this gap, this study aims to measure the association between bullying victimization on cortical and subcortical brain development using structural MRI data collected at three time points: ages 14, 19, and 22. Unlike prior work limited to two time points and a small set of predefined regions, our study leverages three MRI time points and a whole-brain approach to capture more nuanced, region-specific trajectories. We hypothesize that increased bullying victimization is associated with widespread variation in brain development, particularly in regions involved in emotion regulation and social processing. Specifically, we expect to observe distinct patterns of volumetric change, such as decreased cortical and increased subcortical volume change across adolescence and early adulthood. In addition, we will conduct exploratory analyses to examine potential sex differences in bullying-related brain development. Emerging evidence suggests that males and females differ in both neurobiological stress responses and developmental trajectories (19, 20). By examining sex differences, we aim to clarify whether the neurodevelopmental associations of bullying victimization vary by sex, contributing to a more nuanced understanding of brain maturation.

## Materials and Methods

### Participants

This study used data from the IMAGEN project, a European multicenter research initiative investigating how various factors influence brain development and mental health in adolescents. For a comprehensive description of the project’s methodology, refer to Schumann et al. (21). Participants were assessed at eight sites across England, Ireland, France, and Germany. Initial data were collected at age 14, with follow-up assessments conducted at ages 16, 19, and 22. This study specifically analyzes data from ages 14, 19, and 22; data from age 16 were excluded, as MRI scans were not conducted at that time point. A detailed overview of recruitment procedures and inclusion/exclusion criteria is presented in eTable 1, with further details available in Schumann et al. (21).

Written informed consent was obtained from participants and their parent/guardian prior to enrollment. The IMAGEN study was approved by ethics committees at each site, including King’s College London, University of Nottingham, Trinity College Dublin, University of Heidelberg, Technische Universität Dresden, Commissariat à l’Energie Atomique et aux Energies Alternatives, and University Medical Center.

### Bullying victimization

Bullying victimization was assessed using items adapted from the revised Olweus Bully/Victim Questionnaire (22) at all three study timepoints (∼14, ∼19, and ∼22 years). Participants responded to four items evaluating the frequency of victimization over the past six months, each rated on a 5-point Likert scale:

1. “I was bullied at school (a student/peer said or did nasty or unpleasant things to me).”
2. “I was called mean names, made fun of, or teased in a hurtful way by a student/peer.”
3. “A student/peer left me out of things on purpose, excluded me from their group of friends, or completely ignored me.”
4. “I was hit, kicked, pushed or shoved around, or locked indoors by a student/peer.”

Response options ranged from 1 (“never”) to 5 (“three or more times a week”). Scores from the four items were summed and standardized (z-scores) to generate bullying victimization scores at ages 14, 19, and 22, with higher scores indicating increased frequency of self-reported bullying victimization. The composite Olweus Bully/Victim Questionnaire score demonstrated excellent internal consistency in the current sample (Cronbach’s α = 0.89).

### MRI Acquisition and Protocol

Structural magnetic resonance imaging (MRI) data were acquired across eight IMAGEN sites in Europe, all using 3T MRI systems (Siemens: 5 sites; Philips: 2 sites; General Electric: 1 site). High-resolution anatomical scans were obtained using a 3D T1-weighted magnetization-prepared rapid gradient echo (MPRAGE) sequence, aligned with the Alzheimer’s Disease Neuroimaging Initiative (ADNI) protocol. Full details of the IMAGEN MRI acquisition and quality control procedures, including scanner standardization protocols, are available in Schumann et al. (21) and subsequent publications (23, 24), and are accessible via the project’s Standard Operating Procedures (https://imagen-europe.org/). In brief, T1-weighted anatomical images were acquired using a 3D MPRAGE sequence (voxel size = 1.1× 1.1 × 1.1 mm^3^; TR = 2300 ms; TE = 2.9 ms). Full acquisition parameters are provided in eTable 2. All structural MRI scans underwent IMAGEN’s centralized quality control procedure, which included manual inspection for artifacts, data quality, and head motion (see Supplemental Material). Following these procedures, 50 scans were excluded (24 at timepoint 1, 17 at timepoint 2, 9 at timepoint 3), resulting in a final dataset of 4,555 structural MRI scans.

All MRI images were processed using FreeSurfer’s *recon-all* pipeline (version 5.3.0)for full cortical reconstruction and subcortical segmentation (25, 26). This automated process included skull stripping, intensity normalization, Talairach transformation, surface reconstruction, topology correction, and anatomical labeling. Brain parcellation was performed using the Desikan-Killiany-Tourville (DKT) atlas, producing 88 cortical and subcortical regions of interest, from which volume metrics were extracted, along with total gray matter volume (27).

Prior to statistical analyses, outliers were identified as values exceeding ±3 standard deviations from the mean (28). These were visually inspected across timepoints to assess consistency. In line with best practices, only values deemed implausible or inconsistent with developmental trajectories were removed to reduce the influence of artefactual data (29).

### Statistical Analysis

Mixed-effects modeling was used to investigate associations between bullying victimization and longitudinal brain development from adolescence to early adulthood. Analyses were conducted in R (version 4.1.1) using the *lme4* package (version 1.1-27.1) (30). Model fitting followed a structured, top-down selection procedure comprising three sequential stages: (1) determining the optimal developmental trajectory for each region of interest (ROI),(2) specifying the random effects structure, and (3) evaluating the fixed effects of bullying victimization. A complete summary of all models tested is provided in eTable 3. Model selection was guided by a combination of the Akaike Information Criterion (AIC) and likelihood ratio tests (LRT). A model was considered to show improved fit if it demonstrated a reduction in AIC greater than 10 and an LRT p-value < 0.05, in line with established criteria for mixed model comparison (31-33). All mixed-effects models were run with mean-centered continuous variables.

### Step 1: Determining the Optimal Brain Region Developmental Trajectory

To characterize normative patterns of brain development, we first evaluated whether a linear or quadratic representation of time (indexed as months since baseline scan) provided the best fit for each brain region of interest (ROI). This analysis was performed separately for all 88 ROI volumes, including cortical and subcortical structures. Consistent with top-down model selection procedures, both models were initially specified with covariates only— sex, pubertal development status (PDS), socioeconomic status (SES), stressful life events (SLE), and intracranial volume (ICV)—excluding any bullying victimization terms at this stage. See supplemental material for a detailed description of covariates. All models included random intercepts and slopes for participants and random intercepts for scan site. In line with best practices for multisite neuroimaging studies, scan site was modeled as a random effect to account for variance across imaging locations (34). A quadratic time term (time^2^) was retained only when its inclusion significantly improved model fit, allowing for modeling of nonlinear neurodevelopmental trajectories commonly observed during adolescence (18).

### Step 2: Specifying the Random Effects Structure

Following identification of the optimal developmental trajectory, we next determined the most appropriate random effects structure. For both the linear and quadratic models, we compared random intercepts-only models (R1) with models that also included random slopes for time (R2). Scan site was consistently modeled as a random intercept to adjust for multisite acquisition variability. These comparisons were evaluated using the same AIC and LRT thresholds as in Step 1. The selected structure allowed for adequate modeling of within-subject variation in brain development.

### Step 3: Testing Fixed Effects of Bullying Victimization

After finalizing the model structure, we evaluated the fixed effects of bullying victimization using a series of nested models. The null model (F0) included all covariates and time terms but excluded any bullying-related predictors. The simple effects model (F1) added the main effect of bullying victimization frequency, while the interaction model (F2) further included the bullying victimization score × time term to examine whether the association between bullying and brain development varied over time. All models were estimated using maximum likelihood (ML) for model fit comparison, and final parameter estimates were derived using restricted maximum likelihood (REML), which provides unbiased estimates of variance components by accounting for the loss of degrees of freedom when estimating fixed effects (30).

An unstructured variance-covariance matrix was used for random effects, permitting unrestricted estimation of correlations between intercepts and slopes and thus accommodating individual heterogeneity in developmental trajectories (35). To control for multiple comparisons, we applied a false discovery rate (FDR) correction at q < 0.05 using the *stats* package in R (version 4.1.1) (36, 37). This correction was applied to all p-values associated with the bullying victimization score main effects and bullying victimization score × time interactions in the optimally fitted models. Only results that survived FDR correction are reported.

### Sex-Specific Associations Between Bullying Victimization and Brain Development

Exploratory sex-specific analyses were conducted using the optimal models identified during the model selection procedure described above. To test for sex-dependent effects, additional interaction terms for bullying victimization score × sex and bullying victimization score × time × sex were added to these models (F3). As in previous steps, model fit was evaluated using AIC and LRT, and FDR correction was applied to p-values associated with both interaction terms. Only associations that remained statistically significant after correction are reported. These analyses aimed to explore potential sex differences in neurodevelopmental responses to bullying victimization during adolescence.

## Results

### Descriptive Statistics

Full demographic and clinical characteristics are presented in Table 1. Of the 4,555 available MRI scans, 43 were excluded due to missing bullying questionnaire data (Time 1 =11, Time 2 = 30, Time 3 = 2). The final analytic sample comprised 4,512 scans from 2,094 participants (females: 2,171; males: 2,341), spanning an age range of 13.23 to 25.11 years across three time points (see eFigure 1). Bullying victimization showed modest but statistically significant rank-order stability across waves (T1–T2: ρ = 0.24, p < .000001; T2–T3: ρ = 0.20, p < .000001; T1–T3: ρ = 0.19, p < .000001). Attrition analyses were conducted using baseline data and are reported in the Supplementary Material.

**Table 1:** Demographic and Site Characteristics Across Three Time Points.

| Variable Name | Time 1 | Time 2 | Time 3 |
| --- | --- | --- | --- |
| N | 2052 | 1328 | 1132 |
| Sex (female %) | 1009 (49.17%) | 632 (47.59%) | 530 (46.81%) |
| Age | 14.44 (0.39) | 19.01 (0.78) | 22.56 (0.65) |
| OB/VQ score (range) | 4-20 | 0-16 | 0-16 |
| SES (mean) | 0.71 (1.09) | - | - |
| Pubertal Status (mean) | 3.60 (0.71) | - | - |
| Ethnicity (White %) | 89.4% | - | - |
| **Number of Scans at IMAGEN Site** |  |  |  |
| London | 251 | 168 | 153 |
| Northampton | 343 | 211 | 169 |
| Dublin | 192 | 122 | 116 |
| Berlin | 252 | 118 | 155 |
| Hamburg | 261 | 181 | 153 |
| Mannheim | 245 | 155 | 140 |
| Paris | 253 | 204 | 124 |
| Dresden | 255 | 169 | 122 |

**Legend:** N = total number of participants assessed at each respective time point. The Olweus Bully/Victim Questionnaire (OB/VQ) is used to measure bullying victimization and bullying experiences. Pubertal status was assessed via self-report using the Pubertal Development Scale (PDS). Socioeconomic status (SES) was indexed using the family stresses subsection of the Development and Well-Being Assessment (DAWBA). Data collection for Time 1 occurred between 2008–2010, for Time 2 between 2013–2015, and for Time 3 between 2016–2019.

### Bullying Victimization and Brain Development

Model selection fit statistics are presented in the supplemental material (eTables 1–5), with the final models derived from the top-down selection procedure shown in eTable 6. Full model results are provided in eTables 7–10 and eFigure 2.

Significant associations between bullying victimization scores and brain volume development were identified in 30 regions, indicating consistent structural differences across adolescence and early adulthood (Figures 1 and 2). These included widespread volume reductions in cortical areas such as the orbitofrontal cortex, superior and rostral middle frontal gyri, precuneus, precentral and postcentral gyri, insula, entorhinal cortex, temporal pole, and regions within the parietal, occipital, and cerebellar cortices. In contrast, increased volumes were observed in several limbic and subcortical structures, including the amygdala, hippocampus, parahippocampal gyrus, putamen, caudate, nucleus accumbens, and frontal pole. Additional volume reductions were also found in the thalamus, pallidum, and ventral diencephalon.

**Figure 1:**
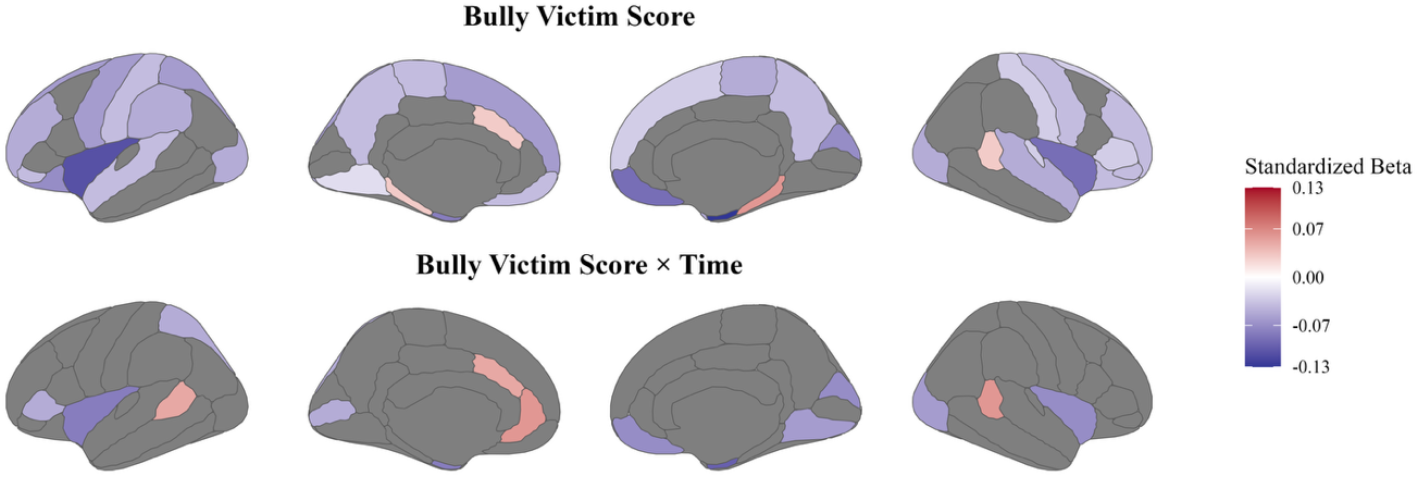
Magnitude of Effect Sizes for Bullying victimization and Bullying victimization-by-Age on Cortical Brain Regions. **Figure 1 legend:** This figure presents standardized effect sizes (β) for the associations between 614 bullying victimization score and the bullying victimization score-by-time interaction on 615 cortical brain volumes. The top panel displays the effect sizes for the main effect of bullying 616 victimization scores, while the bottom panel illustrates the effect sizes for the interaction 617 between bullying victimization and time. The color scale represents standardized β values, with 618 warmer shades indicating stronger positive or negative effects. These visualizations highlight 619 cortical regions where bullying victimization and its interaction with time are significantly 620 linked to changes in brain volume development.

**Figure 2:**
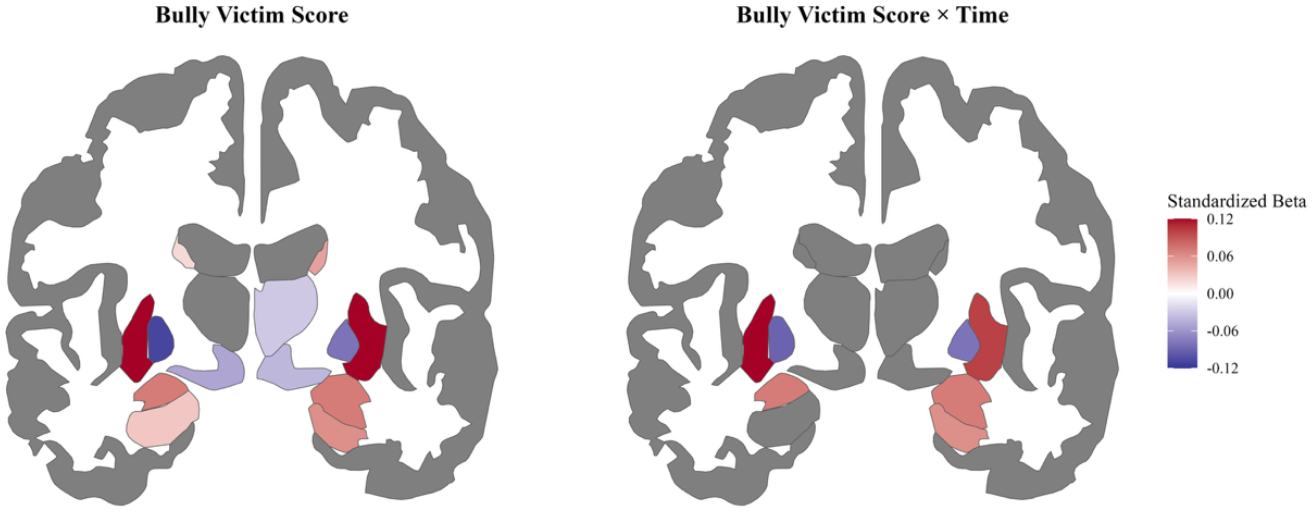
Magnitude of Effect Sizes for Bullying victimization and Bullying victimization-by-Age on Subcortical Brain Regions. **Figure 2 legend:** This figure displays standardized effect sizes (β) for the associations between bullying victimization score and bullying victimization score-by-time on subcortical brain volumes. The left panel illustrates the effect sizes for the main effect of bullying victimization scores, while the right panel shows the effect sizes for the interaction between bullying victimization and time. The color scale reflects standardized β values, with warmer colors indicating larger effect sizes in either the positive or negative direction. These visualizations highlight subcortical regions where bullying victimization and its interaction with time are significantly associated with volume development.

Significant bullying victimization-by-time interactions were identified in approximately 16 regions, indicating that bullying victimization frequency was associated with altered neurodevelopmental trajectories (Figures 1 and 2). Higher levels of victimization frequency were linked to accelerated volumetric growth in regions such as the amygdala, hippocampus, putamen, anterior cingulate cortex, and banks of the superior temporal sulcus. Conversely, reduced volumetric growth was observed in the insula, entorhinal cortex, cerebellum, and visual and parietal cortices (e.g., cuneus, lingual gyrus, superior parietal lobule). These findings suggest that bullying may influence not only regional brain structure but also the pace of brain maturation from adolescence into early adulthood.

### Sex Differences in the Associations of Bullying Victimization on Brain Development

Exploratory analyses revealed significant three-way interactions between bullying victimization, sex, and time on brain volume development (eTables 11–12). These findings indicate that the association between bullying victimization and age-related brain volume trajectories varies by sex (Figure 3).

**Figure 3:**
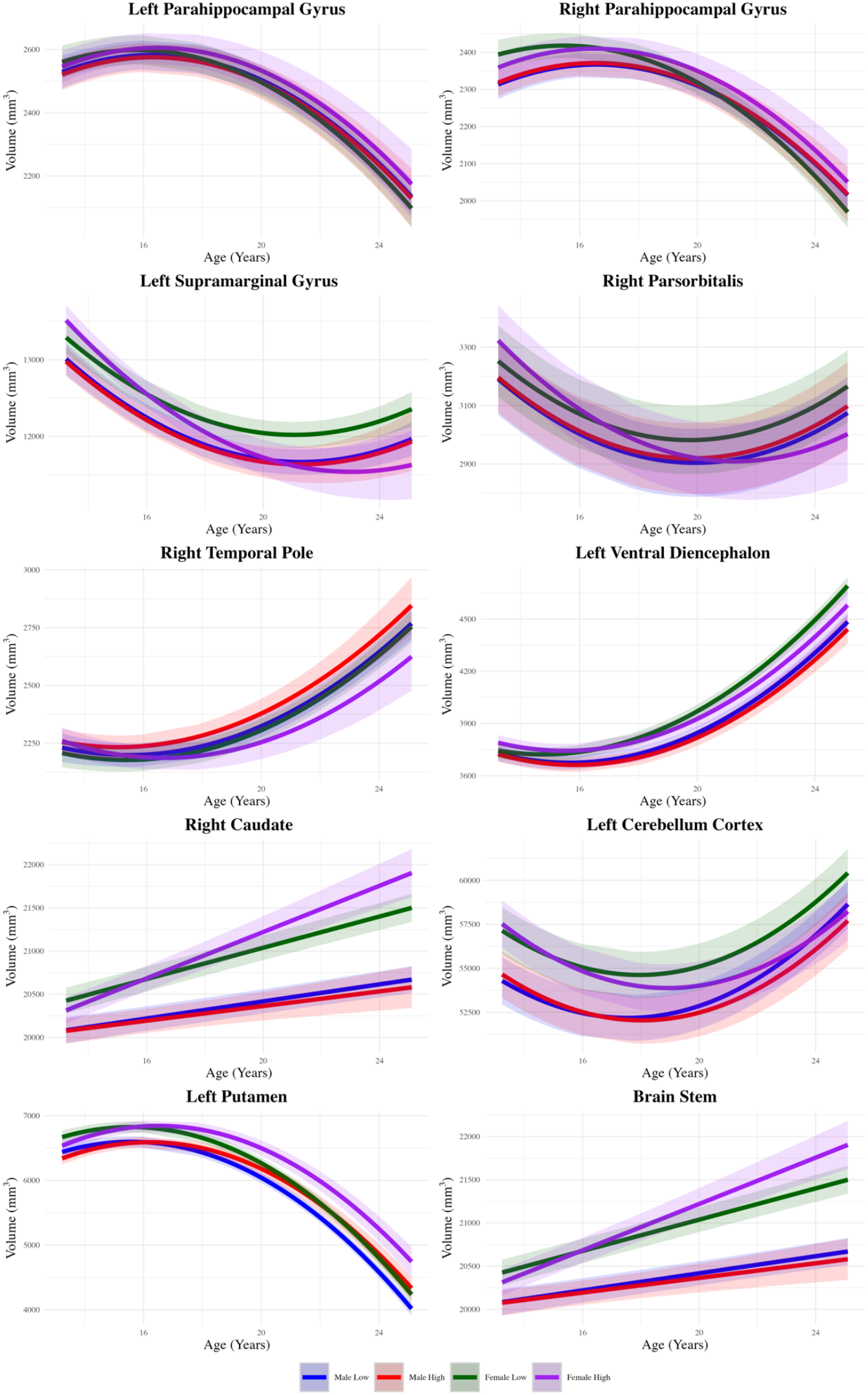
Sex Differences in the Impact of Peer Victimization on Brain Development. **Figure 3 legend:** The analysis had peer victimization as a sum score, but for visualization purposes, these scores were grouped into four categories. “Male Low” represents males in the lowest quartile of bullying victim scores, shown in red. “Male High” represents males in the highest quartile of bullying victim scores, shown in blue. “Female Low” represents females in the lowest quartile of bullying victim scores, shown in dark green. “Female High” represents females in the highest quartile of bullying victim scores, shown in purple. The shaded region around each line represents the 95% confidence interval for the predicted values.

Higher bullying victimization frequency over time was linked to greater volume increases in females compared to males in several subcortical and limbic regions, including the bilateral parahippocampal gyrus, right caudate, left putamen, and brainstem. Conversely, males showed relatively greater volume increases in regions such as the left supramarginal gyrus, right pars orbitalis, right temporal pole, left ventral diencephalon, and left cerebellar cortex.

These results point to sex-specific patterns in the association between bullying victimization and brain development, with distinct regional trajectories emerging across adolescence and early adulthood.

## Discussion

This three-time-point longitudinal neuroimaging study of bully victimization examined the association between bullying victimization and brain development across adolescence and early adulthood. The principal findings are: (1) bullying victimization frequency is significantly associated with altered development in a wide range of cortical and subcortical brain structures, and (2) these alterations show notable sex-specific patterns.

The novelty of this study lies in capturing widespread non-linear developmental brain changes not observed in previous studies, highlighting that the neurodevelopmental associations of bullying victimization may be more widespread than previously known (15, 16). These findings are robust across variables such as sex, age, MRI scanner site, socioeconomic status, pubertal status, and other negative life events, suggesting that the observed brain differences may be uniquely attributed to bullying victimization.

### Association Between Bullying Victimization and Brain Development

#### Habit formation, emotional salience and stress regulation: striatal and subcortical systems

Subcortical structures appear to be particularly sensitive to bullying victimization, with significant volumetric increases observed in the structures of the basal ganglia, including the caudate, putamen, and nucleus accumbens. These regions are integral to motor control, emotional regulation, and reward processing (38). The observed differences in the volumes of the caudate and putamen corroborate previous findings (16). Meanwhile, the discovery of increased volumes in the nucleus accumbens and pallidum represents novel contributions to the field.

Prolonged bullying may trigger neuroplastic adaptations as the brain attempts to cope. Enlarged dorsal striatal structures (caudate and putamen), involved in automatic responses and attention, may underlie increased striatum-dependent (“habit”) learning in bullied individuals (5). These individuals often rely on coping behaviors shaped by past threats, which may be maladaptive in safe contexts, contributing to social distress and difficulty adapting to new environments. Victimized adolescents also show a shift toward striatal-dependent memory processing, associated with cognitive inflexibility and anxiety (39). This type of memory primarily involves the ventral striatum (nucleus accumbens), which plays a central role in negative emotional processing and is linked to internalizing and externalizing symptoms in bullied youth (40). Thus, its enlargement may reflect a heightened bias toward emotionally salient memory encoding. Though less studied, the pallidum also shows volumetric disruptions in this context and has been associated with depression, anxiety, and OCD (41-43). As part of the cortico-striato-thalamo-cortical (CSTC) circuitry, the pallidum is critical for emotional regulation, and its enlargement may reflect CSTC disruption, contributing to stress sensitivity and emotional dysregulation (41, 44).

A novel finding, that contradicts our hypothesis was the association between increased bullying exposure and reduced ventral diencephalon volume. This region, which includes the hypothalamus, is essential for regulating the neuroendocrine stress response (45). Reduced volume here may impair hormonal regulation, compromising the body’s ability to manage stress effectively (46, 47), and increasing susceptibility to anxiety, mood disorders, and other stress-related conditions (48).

#### Emotional reactivity and memory biases: limbic regions

Significant differences were also observed in the limbic system, particularly the amygdala and hippocampus. Enlarged hippocampal volume aligns with prior findings (15), while increased amygdala volume was not previously reported, possibly due to methodological differences such as the binary classification of victimization status in earlier studies. It is plausible that amygdala enlargement reflects heightened emotional reactivity to chronic stress (49). Victimized individuals often show increased neural responses to emotional stimuli, suggesting enhanced stress sensitivity (50, 51), which may contribute to elevated risk for anxiety, depression, and related disorders (52, 53).

Hippocampal enlargement may reflect neuroplastic adaptations to prolonged stress, such as increased neurogenesis or dendritic branching in response to emotionally salient memories (45). These neuroplastic changes may underpin the negative emotional memory biases and increased false memory recall observed in individuals who have experienced bullying victimization (46). Specifically, violent and aggressive false memories have been shown to be positively associated with bullying victimization (54). This heightened memory processing demand could help explain the observed structural increases. Despite these findings, mixed results regarding hippocampal volume call for further research to clarify its role and long-term implications in bullying-related neurodevelopment.

#### Executive control, emotional regulation, and social cognition: Frontal, temporal, parietal, and occipital lobes

Alterations were also noted in various cortical areas across the frontal, temporal, parietal, and occipital lobes. In the frontal lobe, changes were observed in regions such as the medial orbitofrontal cortex, superior frontal gyrus, and frontal pole. Previous research has linked functional and structural alterations in these areas to bullying victimization, suggesting that victims may have difficulties in regulating emotions and making decisions under stress (14). Our findings support these previous studies, reinforcing the idea that bullying impacts critical areas involved in executive functioning (5).

The temporal regions, including the superior temporal gyrus and parahippocampal gyrus, showed changes that could affect memory and emotional association. Studies have indicated that victims of bullying often exhibit altered temporal lobe structures, contributing to difficulties in processing and recalling emotionally charged memories (5). Our results are consistent with these findings, suggesting a common pathway through which bullying affects memory and emotional processing (5).

Significant volume reductions were found in the insula, which are novel findings not previously reported in the literature. Prior research has highlighted significant differences in insula activity in the context of bullying victimization, but structural changes had not been documented until now (55). The insula is a deep cortical structure critical for emotional processing and interoceptive awareness (5). Alterations in this structure have been linked to heightened sensitivity to emotional stimuli and social rejection, a common feature of bullying victimization (5).

#### Environmental interpretation via perceptual, predictive, and integrative systems: parietal, occipital and cerebellum regions

Alterations in the parietal and occipital lobes, such as the precuneus and cuneus, may influence how victims process and respond to visual and spatial cues (56, 57). This can impact their social interactions and stress responses (5). Previous studies have found that these regions, when affected by bullying, can alter how individuals perceive and react to their environment, potentially leading to social withdrawal and heightened anxiety (5). Our findings align with these studies, suggesting that visual and spatial processing deficits may contribute to the social challenges faced by bullied individuals (5).

Our study revealed a novel link between bullying victimization frequency and reduced cerebellar volume, suggesting that prolonged social stress may impair cerebellar development. While traditionally associated with motor coordination, the cerebellum also plays a key role in social and cognitive processes essential for social interaction (58). Poor motor skills, governed by cerebellar function, are strong predictors of being bullied (59), as individuals with less refined motor abilities may struggle with social coordination and integration. Beyond motor control, the cerebellum contributes to higher-order functions such as prediction, error-based learning, and emotional recognition (58). Disruptions in these mechanisms may impair interpretation of social cues, leading to negative emotional biases common in bullied individuals, who often misread neutral or ambiguous interactions as hostile, thus increasing their vulnerability to interpersonal difficulties and psychopathology (58, 60).

In summary, our findings indicate that bullying victimization is linked to widespread cortical alterations affecting cognitive and emotional functioning. These structural changes may contribute to the psychological and behavioral difficulties seen in victims and support a neurobiological cycle in which bullying exacerbates vulnerability to further victimization and long-term mental health risks.

### Sex-Specific Differences in Brain Volume Development

This study identified distinct sex-specific patterns of brain volume changes associated with bullying victimization frequency, suggesting that males and females may exhibit different neurodevelopmental adaptations to similar social stressors. Females demonstrated relatively greater volume increases in subcortical and limbic regions—specifically the bilateral parahippocampal cortex, right caudate, left putamen, and brainstem—while males showed increased volume predominantly in a combination of cortical and subcortical regions, including the left supramarginal gyrus, right pars orbitalis, right temporal pole, left ventral diencephalon, and left cerebellar cortex.

These divergent patterns may reflect differences in the nature of bullying typically experienced by each sex. Females are more commonly exposed to relational bullying—such as social exclusion, manipulation, and rumour-spreading (61)—which preferentially engages limbic and paralimbic networks involved in emotional memory and social cognition (62-65). In contrast, males more frequently encounter physical and overt forms of bullying, including verbal aggression and threats, which may more strongly recruit sensorimotor and salience-detection circuits, consistent with volumetric changes observed in regions such as the cerebellum and ventral diencephalon (66-68) .

Together, they underscore the importance of considering sex as a moderating factor in the neural associations of bullying victimization. Although these insights are promising, further research is needed to elucidate the underlying mechanisms and their long-term neurodevelopmental relevance.

### Neurobiological Mechanisms Linked to Brain Volume Alterations in Bullying Victimization

The region-specific brain volume changes associated with bullying victimization likely reflect broader neurobiological mechanisms triggered by chronic social stress. One key system implicated is the hypothalamic–pituitary–adrenal (HPA) axis, which becomes activated in response to psychological stress and elevates glucocorticoid levels—a pattern observed in bullied children (69-72). These glucocorticoids disproportionately affect multiple brain regions, including the prefrontal cortex (PFC) and hippocampus (73). Sustained cortisol exposure can impair synaptic plasticity, promote dendritic atrophy, and even lead to neurodegeneration, which may underlie the reduced PFC volume observed in bullied youth (74). Interestingly, although chronic stress is typically associated with hippocampal atrophy (75), this study and previous research found that bullying victimization is associated with increased hippocampal volume development during early life (15). This may reflect an early adaptive response, where initial stress exposure temporarily enhances dendritic complexity or glial activity before volume declines with prolonged stress (76, 77). Similarly, increased volume in the ventral diencephalon, including the hypothalamus, may reflect stress-induced plasticity within neuroendocrine circuits (78). Chronic stress enhances corticotropin-releasing hormone (CRH) production, increases excitatory input, and reduces inhibitory control in the paraventricular nucleus, sustaining HPA axis activation (78). These adaptations may support short-term homeostasis under prolonged social threat and contribute to the observed volumetric increase.

Beyond the HPA axis, bullying may also activate the locus coeruleus–norepinephrine (LC-NE) system, a brainstem arousal network responsive to threat (79). Sustained norepinephrine release is associated with increased amygdala excitability and synaptic activity (80), which over time may drive structural plasticity—such as dendritic hypertrophy (81)—potentially contributing to the amygdala volume increases observed in bullied adolescents. A concurrent reduction in medial PFC volume—essential for top-down regulation of the LC-NE system—could reflect a shift toward reflexive, emotion-driven processing (82), potentially explaining heightened threat sensitivity in bullying victims (83).

Additionally, volumetric increases in the caudate, putamen, and nucleus accumbens may result from stress-induced changes in dopaminergic signalling within mesostriatal and mesolimbic pathways (84). This dopaminergic hyperactivity has been linked to structural plasticity, including dendritic growth and increased spine density in medium spiny neurons— cellular changes that may underlie the volumetric expansion of striatal regions (85). These changes may reflect an adaptive response to repeated exposure to socially salient stressors, enhancing the salience of threat cues and reinforcing habitual coping behaviors (86). Over time, this plasticity could contribute to inflexible behavioral patterns and heightened sensitivity to social stress observed among individuals exposed to bully victimization (39)

While these findings point to region-specific developmental adaptations, the neurobiological mechanisms underlying altered brain development in bullied youth are inherently complex and likely involve the dynamic interplay of hormonal, neuroimmune, and neurotransmitter systems. Continued investigation into these pathways is essential, as developing a neurobiological framework is critical for understanding the long-term impact of bullying victimization.

### Limitations

This study has several limitations that warrant consideration. Bullying victimization in this study was measured from age 14 onward, potentially missing developmental changes from earlier experiences. As a result, early influences on brain development may be underrepresented, including critical periods affected by prior victimization. Future research should incorporate earlier and broader longitudinal data to better capture these effects. Another limitation of the current study is that we were unable to distinguish between school-based and out-of-school bullying experiences, as participant educational status was not consistently recorded. Future research should explore how the context of bullying may differentially impact developmental trajectories. While the questionnaire included items that could reflect cyberbullying, it was not specifically measured, which may have led to an underestimation of victimization. Future studies should use dedicated cyberbullying measures, such as those by Aricak et al. (87), to better assess its impact on brain development. Ethnicity was not included as a covariate due to imbalanced representation across sites. While scan site served as a proxy-with 89.4% of participants aligning with the dominant local ethnic group - this does not fully capture ethnic variation. More diverse and balanced samples are needed to examine ethnicity’s role in bully victimization and neurodevelopment. As IMAGEN is a community-based cohort, clinical diagnoses of pre-existing mental health conditions were not collected. This makes it difficult to determine whether observed brain changes are driven by bullying victimization alone or influenced by earlier psychiatric vulnerabilities. With only three timepoints per participant, higher-order nonlinear models (e.g., cubic, biquadratic) were not feasible due to the risk of overfitting and model instability (88, 89). Linear and quadratic trajectories, however, are well-established in adolescent brain development (17), making them a statistically appropriate and theoretically grounded choice. The sparse longitudinal sampling also limited our ability to reliably test whether within-person changes in bullying victimization were paralleled by changes in cortical developmental trajectories, which will be an important focus for future studies using denser longitudinal designs better suited to capturing dynamic within-person victimization trajectories.

## Conclusion

This longitudinal MRI study demonstrates that bullying victimization is associated with widespread structural differences in brain development from adolescence to early adulthood, with distinct sex-specific patterns. These findings extend prior work by highlighting neurodevelopmental alterations across circuits involved in stress, emotional learning, and social cognition. While causality cannot be inferred, the results underscore bullying victimization as a salient social experience linked to long-term variation in brain maturation and provide a foundation for future neuroimaging research to explore the underlying mechanisms driving mental health vulnerability.

## Disclosures

Dr Banaschewski served in an advisory or consultancy role for AGB Pharma, eye level, Infectopharm, Medice, Neurim Pharmaceuticals, Oberberg GmbH and Takeda. He received conference support or speaker’s fee by Janssen-Cilag, Medice and Takeda. He received royalities from Hogrefe, Kohlhammer, CIP Medien, Oxford University Press; the present work is unrelated to these relationships. Dr Barker has received honoraria from General Electric Healthcare for teaching on scanner programming courses. Dr Poustka served in an advisory or consultancy role for Roche and Viforpharm and received speaker’s fee by Shire. She received royalties from Hogrefe, Kohlhammer and Schattauer. The present work is unrelated to the above grants and relationships. The other authors report no biomedical financial interests or potential conflicts of interest.

## Acknowledgments

This work received support from the following sources: the European Union-funded FP6 Integrated Project IMAGEN (Reinforcement-related behaviour in normal brain function and psychopathology) (LSHM-CT-2007-037286), the Horizon 2020 funded ERC Advanced Grant ‘STRATIFY’ (Brain network based stratification of reinforcement-related disorders) (695313), Horizon Europe ‘environMENTAL’, grant no: 101057429, UK Research and Innovation (UKRI) Horizon Europe funding guarantee (10041392 and 10038599), Human Brain Project (HBP SGA 2, 785907, and HBP SGA 3, 945539), the Chinese government via the Ministry of Science and Technology (MOST). The German Center for Mental Health (DZPG), the Bundesministerium für Bildung und Forschung (BMBF grants 01GS08152; 01EV0711; Forschungsnetz AERIAL 01EE1406A, 01EE1406B; Forschungsnetz IMAC-Mind 01GL1745B), the Deutsche Forschungsgemeinschaft (DFG project numbers 458317126 [COPE], 186318919 [FOR 1617], 178833530 [SFB 940], 386691645 [NE 1383/14-1],402170461 [TRR 265], 454245598 [IRTG 2773]), the Medical Research Foundation and Medical Research Council (grants MR/R00465X/1 and MR/S020306/1), the National Institutes of Health (NIH) funded ENIGMA-grants 5U54EB020403-05, 1R56AG058854-01 and U54 EB020403 as well as NIH R01DA049238, the National Institutes of Health, Science Foundation Ireland (16/ERCD/3797). NSFC grant 82150710554. Further support was provided by grants from: - the ANR (ANR-12-SAMA-0004, AAPG2019 - GeBra), the Eranet Neuron (AF12-NEUR0008-01 - WM2NA; and ANR-18-NEUR00002-01 - ADORe), the Fondation de France (00081242), the Fondation pour la Recherche Médicale (DPA20140629802), the Mission Interministérielle de Lutte-contre-les-Drogues-et-les-Conduites-Addictives (MILDECA), the Assistance-Publique-Hôpitaux-de-Paris and INSERM (interface grant), Paris Sud University IDEX 2012, the Fondation de l’Avenir (grant AP-RM-17-013), the Fédération pour la Recherche sur le Cerveau.

## Data Availability

All the data are available from the authors upon reasonable request and with permissions of the IMAGEN consortia. https://github.com/imagen2/imagen_mri

## Code Availability

https://github.com/mconnaug/Bullying_Brain_Development

## Notes

### Competing Interest Statement

Dr Banaschewski served in an advisory or consultancy role for Lundbeck Medice Neurim Pharmaceuticals Oberberg GmbH Shire. He received conference support or speakers fee by Lilly Medice Novartis and Shire. He has been involved in clinical trials conducted by Shire and Viforpharma. He received royalties from Hogrefe Kohlhammer CIP Medien Oxford University Press. The present work is unrelated to the above grants and relationships. Dr Barker has received honoraria from General Electric Healthcare for teaching on scanner programming courses. Dr Poustka served in an advisory or consultancy role for Roche and Viforpharm and received speakers fee by Shire. She received royalties from Hogrefe Kohlhammer and Schattauer. The present work is unrelated to the above grants and relationships. The other authors report no biomedical financial interests or potential conflicts of interest.

### Summary of Updates

Additional sensitivity analyses, and updated figures and tables have been included to increase robustness of the studies findings.

