## Supplemental Material for "Bullying Victimization and Brain Development: A Longitudinal Structural Magnetic Resonance Imaging Study from Adolescence to Early Adulthood"

*Connaughton et al.,*

### 1 Methods

#### *1.1* IMAGEN study inclusion/exclusion criteria

Recruitment procedures, detailed inclusion and exclusion criteria, and Standard Operating Procedures for the IMAGEN study have been previously described by Schumann et al. (2010) (1). Briefly, participants were recruited based on the following criteria:

**eTable 1:** Summary of IMAGEN study Inclusion and Exclusion Criteria.

| Category | Criterion | Action |
|---|---|---|
| **A. Demographics** | Child aged 14 years | Inclusion |
| **B. Pregnancy and Birth History** | Maternal alcohol consumption exceeding 210 ml per week during pregnancy (equivalent to approximately 14 beers, 9 glasses of wine, or 7 servings of spirits) | Exclusion |
| | Maternal diabetes diagnosed before pregnancy and treated with insulin | Exclusion |
| | Premature birth (< 35 weeks' gestation) or history of placental abruption | Exclusion |
| | Severe hyperbilirubinemia requiring transfusion | Exclusion |
| **C. Child's Medical History** | Diagnosis of type 1 diabetes | Exclusion |
| | Systemic rheumatologic diseases (e.g., glomerulonephritis, endocarditis secondary to streptococcal infection) | Exclusion |
| | History of malignant tumors requiring chemotherapy (e.g., leukemia) | Exclusion |
| | Congenital heart defects or history of heart surgery | Exclusion |
| | Presence of aneurysms | Exclusion |
| **D. Neurological Conditions** | Diagnosis of epilepsy | Exclusion |
| | History of bacterial infections affecting the central nervous system | Exclusion |
| | History of brain tumors | Exclusion |
| | Head injury involving loss of consciousness lasting longer than 30 minutes | Exclusion |
| | Diagnosis of muscular dystrophy or myotonic dystrophy | Exclusion |
| | Nutritional or metabolic diseases (e.g., phenylketonuria, failure to thrive) | Exclusion |
| **E. Developmental and Sensory Disorders** | Major neurodevelopmental disorders (e.g., autism spectrum disorders) | Exclusion |
| | Hearing impairment requiring the use of hearing aids | Exclusion |

| | | |
|---|---|---|
| | Vision disorders not correctable with standard treatments (e.g., strabismus, significant visual deficits) | Exclusion |
| **F. Psychiatric and Cognitive Status** | Ongoing treatment for schizophrenia or bipolar disorder | Exclusion |
| | Intelligence quotient (IQ) below 70 | Exclusion |
| | Presence of metallic implants | Exclusion |
| **G. MRI Contraindications** | Presence of electronic implants (e.g., pacemakers) | Exclusion |
| | Severe claustrophobia | Exclusion |

#### 1.2 MRI Acquisition Parameters for Structural Imaging (MPRAGE Sequence)

| **eTable 2:** IMAGEN Structural MRI Acquisition Parameters | |
|---|---|
| Sequence Parameter | Value |
| Repetition Time (TR, ms) | 2300 |
| Echo Time (TE, ms) | 2.8 |
| Echo Train Length (ETL) | - |
| Parallel Imaging / Acceleration Factor | N |
| Number of Signal Averages (NSA) | 1 |
| Scan Duration | ~9:20 minutes |
| Excitation Flip Angle (degrees) | 8-9 |
| Acquisition Type | 3D |
| Matrix Size (Frequency Direction) | 256 |
| Matrix Size (Phase Direction) | 256 |
| No. of Slices / Matrix Size (3D) | 160–170 |
| Field of View (FOV) Frequency (cm) | 28.0 |
| Field of View (FOV) Phase (%) | 94% |
| Slice Thickness (mm) | 1.1 |
| Slice Gap (mm) | Not applicable |
| Slice Orientation | Sagittal |
| In-plane Phase Encoding Direction | - |
| Slice Acquisition Order | Not applicable |
| Slice Acquisition Direction | Left → Right |
| Sequence-Specific Parameters | Inversion Time (TI) = 900 ms |
| Legend: MPRAGE = Magnetization-Prepared Rapid Gradient-Echo sequence; TR = Repetition Time; TE = Echo Time; ETL = Echo Train Length; NSA = Number of Signal Averages; FOV = Field of View; TI = Inversion Time; mm = millimetres; ms = milliseconds; N = No parallel imaging applied. All parameters were harmonized according to the Alzheimer's Disease Neuroimaging Initiative (ADNI) standards to ensure comparability across sites. | |

#### 1.3 Distribution imaging assessments by age and sex

**eFigure 1:** Distribution of sample by age and sex.

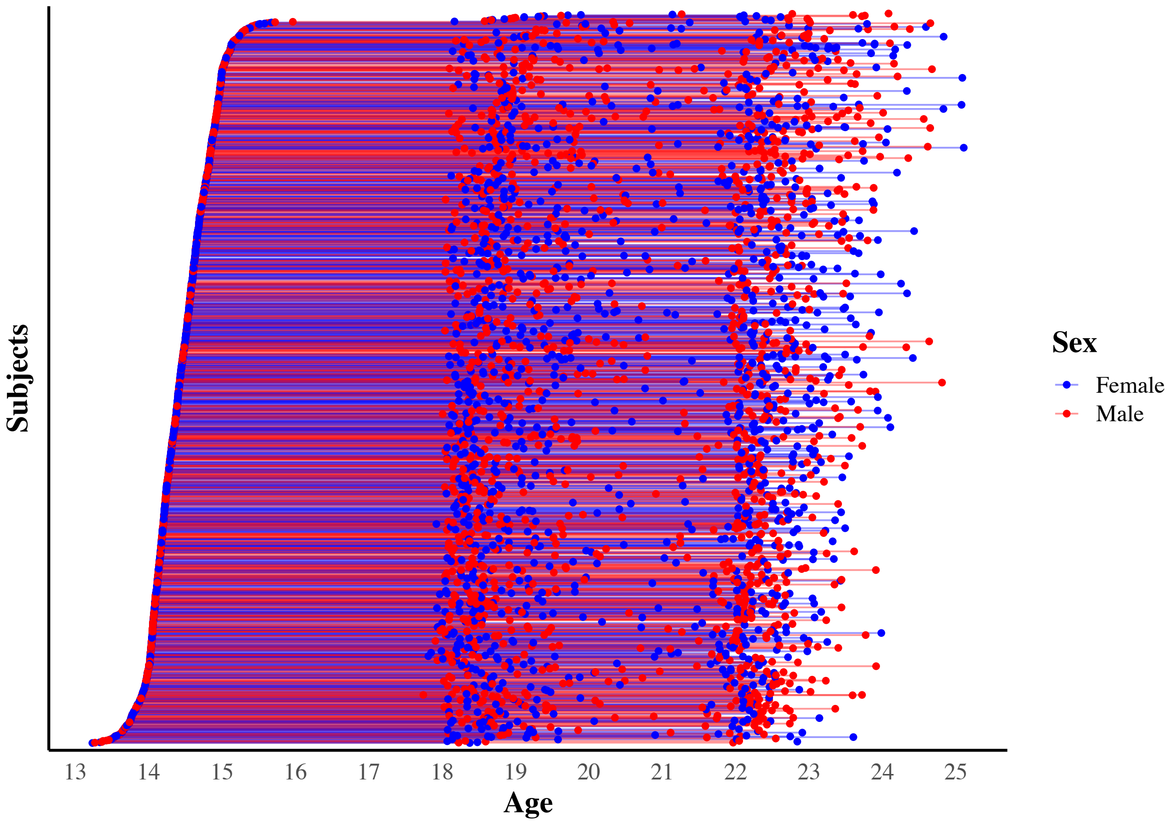


**Legend:** Distribution of MRI scans by age and sex in the study sample, showing the number of scans for each age group by sex

#### 1.4 Quality control procedures

The IMAGEN study has established specific protocols and quality control procedures to ensure the reliability and integrity of clinical, behavioral, neuropsychological, and MRI data. These assessments and quality control processes have been detailed in prior publications from the Consortium (1, 2). The procedures combine automated systems, manual review, and standardized reporting to maintain consistently high data quality across all participating study centers.

##### Clinical, Behavioral, and Neuropsychological Assessments

- Participant feedback on tests administered through the 'Psytools' platform is automatically recorded in the database.

- Research Assistant (RA) quality assessments and comments are documented immediately after each session and manually reviewed.
- Cases with missing reliability ratings or flagged as doubtful are followed up with the respective study centers for clarification or additional information.
- Data judged to be unreliable are excluded from subsequent analyses.
- Behavioral data are systematically checked for outliers, missing values, and deviations from a normal distribution.

**MRI Data**

- Automated and visual (web-based) quality control methods are applied to preprocessed structural and functional MRI data. Scans are flagged for issues such as normalization errors, segmentation problems, clinical abnormalities, motion artifacts, deformations, and susceptibility artifacts.
- Contrast maps are reviewed for outliers and missing values.
- MRI quality reports generated by RAs immediately after scanning are uploaded to the central database and manually reviewed.
- Behavioral log files from MRI tasks are examined for missing, incomplete, or anomalous data points.

The Python script used for these quality control procedures is available at: https://github.com/PhilipSpechler/Visual_QC_for_MRI_Datasets.

#### 1.5 Tested Mixed Effects Models

| **eTable 3:** Mixed models tested: Bullying and Early Brain Development from Adolescence to Early Adulthood. | |
|---|---|
| **Step 1: Determining the Optimal Brain Region Developmental Trajectory** | |
| Linear | ROI ~ OB/VQ*time + sex + SLE + SES + PDS + ICV + (1 + time \|subject) + (1\|scan site) |
| Quadratic | ROI ~ OB/VQ*time + $time^2$ + sex + SLE + SES + PDS + ICV + (1 + time \|subject) + (1\|scan site) |
| **Step 2: Specifying the Random Effects Structure** | |
| Linear-R1 | ROI ~ OB/VQ*time + sex + SLE + SES + PDS + ICV + (1 \|subject) + (1\|scan site) |
| Linear -R2 | ROI ~ OB/VQ*time + sex + SLE + SES + PDS + ICV + (1 + time \|subject) + (1\|scan site) |
| Quadratic-R1 | ROI ~ OB/VQ*time + $time^2$ + sex + SLE + SES + PDS + ICV + (1 \|subject) + (1\|scan site) |
| Quadratic-R2 | ROI ~ OB/VQ*time + $time^2$ + sex + SLE + SES + PDS + ICV + (1 + time \|subject) + (1\|scan site) |
| **Step 3: Testing Fixed Effects of Bullying Victimization** | |

| | |
|---|---|
| Linear-R1-F0 | ROI ~ time + sex + SLE + SES + PDS + ICV + (1 \|subject) + (1\|scan site) |
| Linear-R2-F0 | ROI ~ time + sex + SLE + SES + PDS + ICV + (1 + time \|subject) + (1\|scan site) |
| Quadratic-R1-F0 | ROI ~ time + time$^2$ + sex + SLE + SES + PDS + ICV + (1 \|subject) + (1\|scan site) |
| Quadratic-R2-F0 | ROI ~ time + time$^2$ + sex + SLE + SES + PDS + ICV + (1 + time \|subject) + (1\|scan site) |
| Linear-R1-F1 | ROI ~ OB/VQ + time + sex + SLE + SES + PDS + ICV + (1 \|subject) + (1\|scan site) |
| Linear-R2-F1 | ROI ~ OB/VQ + time + sex + SLE + SES + PDS + ICV + (1 + time \|subject) + (1\|scan site) |
| Quadratic-R1-F1 | ROI ~ OB/VQ + time + time$^2$ + sex + SLE + SES + PDS + ICV + (1 \|subject) + (1\|scan site) |
| Quadratic-R2-F1 | ROI ~ OB/VQ + time + time$^2$ + sex + SLE + SES + PDS + ICV + (1 + time \|subject) + (1\|scan site) |
| Linear-R1-F2 | ROI ~ OB/VQ*time + sex + SLE + SES + PDS + ICV + (1 \|subject) + (1\|scan site) |
| Linear-R2-F2 | ROI ~ OB/VQ*time + sex + SLE + SES + PDS + ICV + (1 + time \|subject) + (1\|scan site) |
| Quadratic-R1-F2 | ROI ~ OB/VQ*time + time$^2$ + sex + SLE + SES + PDS + ICV + (1 \|subject) + (1\|scan site) |
| Quadratic-R2-F2 | ROI ~ OB/VQ*time + time$^2$ + sex + SLE + SES + PDS + ICV + (1 + time \|subject) + (1\|scan site) |
| **Exploratory Analyses: Sex Differences in the Association Between Bullying Victimization and Brain Development** | |
| Linear-R1-F3 | ROI ~ OB/VQ*time*sex + SLE + SES + PDS + ICV + (1 \|subject) + (1\|scan site) |
| Linear-R2-F3 | ROI ~ OB/VQ*time*sex + SLE + SES + PDS + ICV + (1 + time \|subject) + (1\|scan site) |
| Quadratic-R1-F3 | ROI ~ OB/VQ*time*sex + time$^2$ + SLE + SES + PDS + ICV + (1 \|subject) + (1\|scan site) |
| Quadratic-R2-F3 | ROI ~ OB/VQ*time*sex + time$^2$ + SLE + SES + PDS + ICV + (1 + time \|subject) + (1\|scan site) |
| **Legend:** R1 = random intercepts only, R2 = random intercepts and slopes; F0 = null model (no OB/VQ term), F1 = simple fixed effects (main effect of OB/VQ), F2 = complex fixed effects (OB/VQ × time interaction), F3 = exploratory fixed effects (OB/VQ × time × sex interaction); OB/VQ = Olweus Bully/Victim Questionnaire; time = time from baseline scan (in months); SLE = stressful life events; PDS = puberty developmental scale; SES = socioeconomic status; ICV = intracranial volume. | |

#### 1.6 Mixed Effects Model Covariates

To isolate the longitudinal impact of bullying on brain development, we included a set of theoretically and empirically grounded covariates in all mixed-effects models. These covariates included: time, time$^2$ (for quadratic models), intracranial volume (ICV), sex, socioeconomic status (SES), pubertal status, and stressful life events.

As recommended by *Freesurfer* longitudinal analysis, time was modelled as months since the baseline scan, which enables modelling of continuous, age-related brain maturation across adolescence while accounting for variation in follow-up intervals. This approach also defines the model intercept at each participant's baseline (time = 0), improves interpretability of change trajectories, and cleanly separates within-person developmental change from between-person differences in baseline age (3). For quadratic fits, a squared time term (*time*$^2$) was included to allow for non-linear developmental trajectories. This is recommended because many brain structures follow non-linear patterns of change during adolescence, such as accelerated growth followed by plateauing or decline, and including *time²* allows the model to capture these biologically plausible patterns more accurately (3).

ICV was estimated using *FreeSurfer* and included to adjust for variability in overall brain size. ICV is a standard covariate in structural neuroimaging to ensure that regional brain differences reflect relative changes rather than differences in global head size (4). Biological sex (male, female) was also included as a covariate, in line with best practices for MRI studies. Sex differences are known to influence brain structure and development, and adjusting for sex helps improve the specificity and generalizability of findings(4).

SES was indexed using the family stresses subsection of the DAWBA, including indicators such as parental education and employment, financial difficulties, and housing conditions (5). A cumulative score was computed to reflect socioeconomic adversity. Lower SES has been linked to altered trajectories of brain development in adolescence (6) and may increase both vulnerability to bullying and stress-related neurodevelopmental effects (7).

Pubertal status was assessed using the pubertal developmental scale, a 8-item self-report measure based on Tanner staging (8), assessing pubertal markers such as growth, pubic hair, voice changes (males), and menarche (females). In-line with prior longitudinal MRI research (9) pubertal status scores were mode-centred to account for sex differences in the timing of puberty and to provide a meaningful reference point corresponding to the most commonly reported stage in the sample.

Stressful life events (SLE) were assessed using the self-report Life Events Questionnaire (LEQ; (10)), focusing on distressing events (e.g., bereavement, family conflict) rated as negative by participants. Following previous work in the IMAGEN cohort (11, 12), twenty events were identified as stressful based on participant ratings, where events marked as “unhappy” or “very unhappy” were considered emotionally negative. A cumulative stressful life events (SLE) score was calculated by summing the number of these negatively rated events reported by each participant. At age 14, the SLE score reflected events from the past 12 months, whereas at ages 16 and 19, it captured events occurring since the last study visit.

### 2 <u>Results</u>

#### 2.1 Attrition Analyses

Attrition analyses were conducted using baseline (age 14) data only, with one observation per participant. Participants were classified as retained if they completed the final follow-up assessment at age 22 (T3) and non-retained if they contributed baseline data only. Of the 2,063 participants with baseline MRI data, 1,109 (53.8%) were retained to T3 and 954 (46.2%) were non-retained.

Sex distribution differed modestly between groups, with a higher proportion of males in the retained group compared to the non-retained group ($\chi^2(1) = 4.51$, $p = .034$). Retained participants had lower baseline socioeconomic status (SES) than non-retained participants ($t(1789.1) = 5.38$, $p < .001$) and showed slightly more advanced pubertal development at baseline ($t(2031) = -2.81$, $p = .005$).

In contrast, retained and non-retained participants did not differ in baseline bullying victimization frequency ($t(1937.8) = 1.48$, $p = .139$) or estimated total intracranial volume ($t(2044.2) = -0.42$, $p = .676$).

#### 2.2 Bullying Victimization and Brain Development

**eFigure 2:** The Impact of Peer Victimization on Brain Development from Adolescence to Early Adulthood.

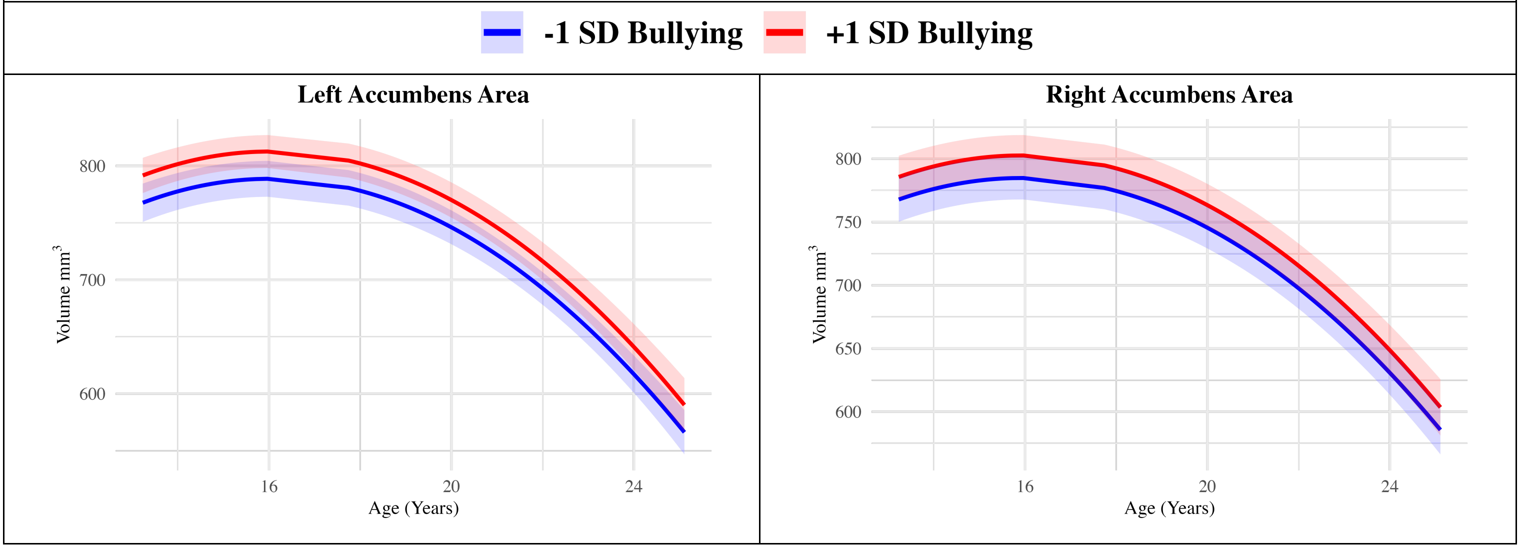

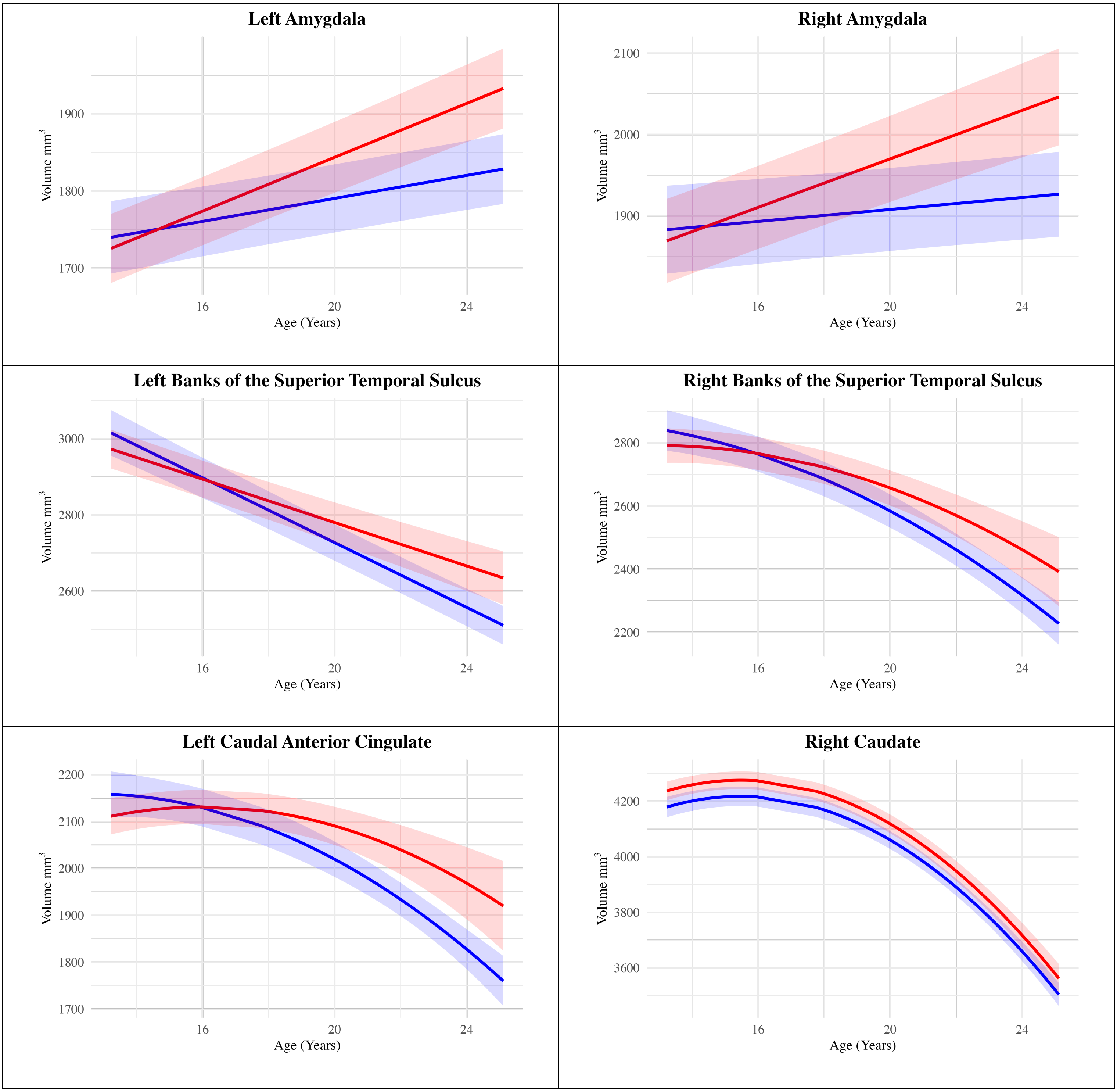
Left Amygdala
Volume mm3
1700
1800
1900
16
20
24
Age (Years)
Right Amygdala
Volume mm3
1900
2000
2100
16
20
24
Age (Years)
Left Banks of the Superior Temporal Sulcus
Volume mm3
2600
2800
3000
16
20
24
Age (Years)
Right Banks of the Superior Temporal Sulcus
Volume mm3
2200
2400
2600
2800
16
20
24
Age (Years)
Left Caudal Anterior Cingulate
Volume mm3
1700
1800
1900
2000
2100
2200
16
20
24
Age (Years)
Right Caudate
Volume mm3
3600
3800
4000
4200
16
20
24
Age (Years)

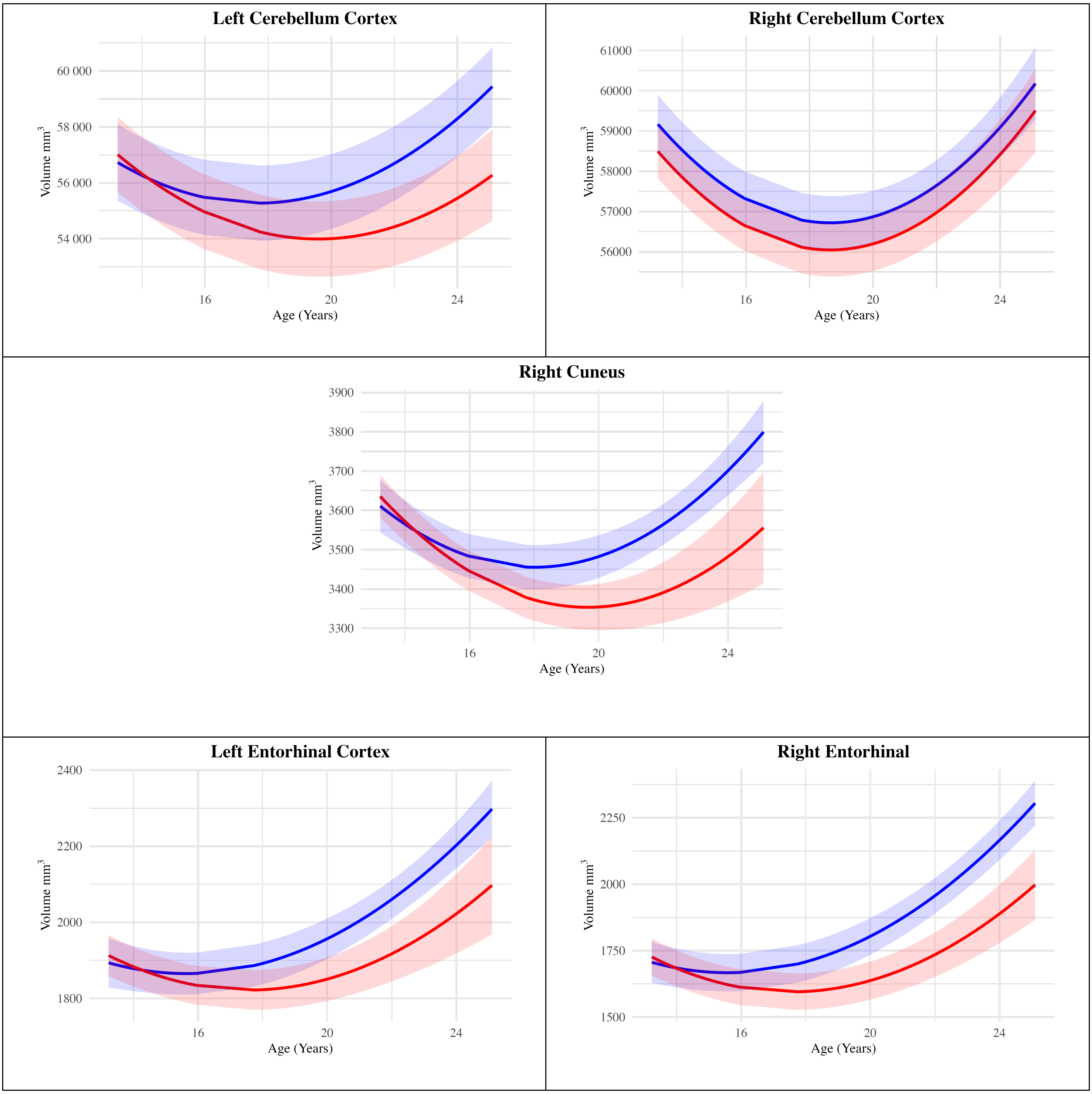
Left Cerebellum Cortex
60 000
58 000
56 000
54 000
Volume mm3
16
20
24
Age (Years)
Right Cerebellum Cortex
61000
60000
59000
58000
57000
56000
Volume mm3
16
20
24
Age (Years)
Right Cuneus
3900
3800
3700
3600
3500
3400
3300
Volume mm3
16
20
24
Age (Years)
Left Entorhinal Cortex
2400
2200
2000
1800
Volume mm3
16
20
24
Age (Years)
Right Entorhinal
2250
2000
1750
1500
Volume mm3
16
20
24
Age (Years)

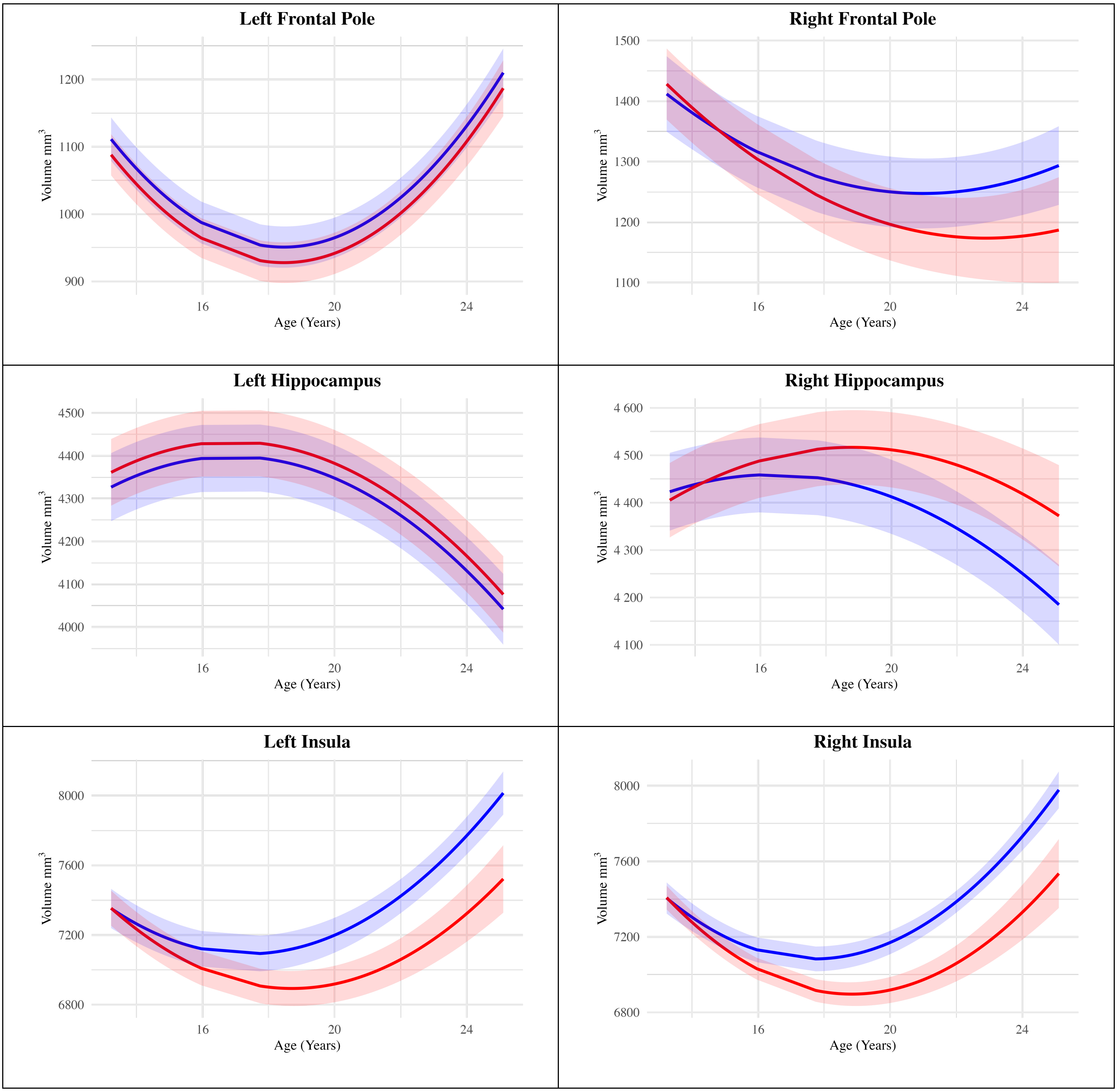
Left Frontal Pole
1200
1100
1000
900
Volume mm3
16
20
24
Age (Years)
Right Frontal Pole
1500
1400
1300
1200
1100
Volume mm3
16
20
24
Age (Years)
Left Hippocampus
4500
4400
4300
4200
4100
4000
Volume mm3
16
20
24
Age (Years)
Right Hippocampus
4 600
4 500
4 400
4 300
4 200
4 100
Volume mm3
16
20
24
Age (Years)
Left Insula
8000
7600
7200
6800
Volume mm3
16
20
24
Age (Years)
Right Insula
8000
7600
7200
6800
Volume mm3
16
20
24
Age (Years)

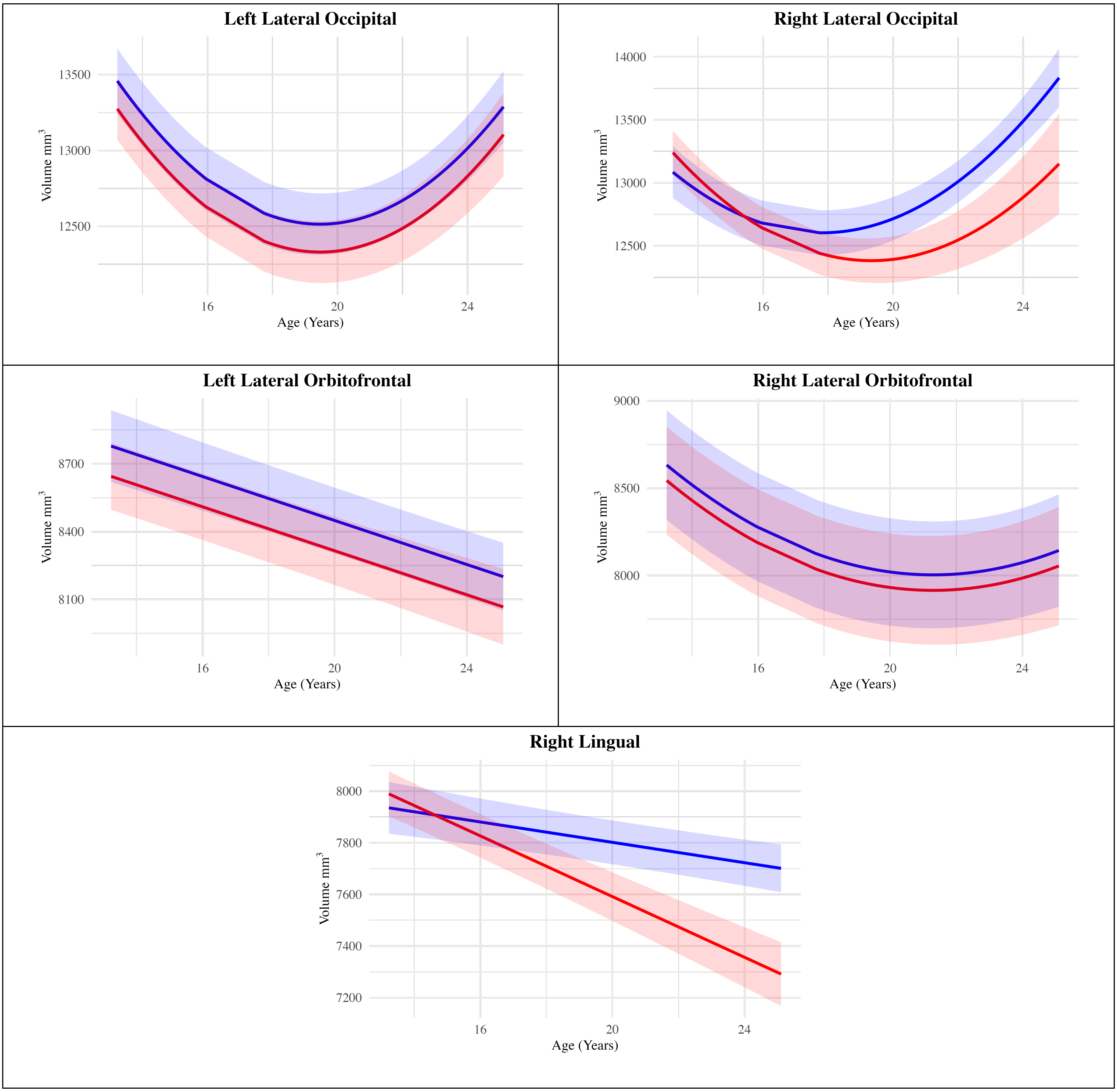
Left Lateral Occipital
13500
13000
12500
Volume mm3
16
20
24
Age (Years)
Right Lateral Occipital
14000
13500
13000
12500
Volume mm3
16
20
24
Age (Years)
Left Lateral Orbitofrontal
8700
8400
8100
Volume mm3
16
20
24
Age (Years)
Right Lateral Orbitofrontal
9000
8500
8000
Volume mm3
16
20
24
Age (Years)
Right Lingual
8000
7800
7600
7400
7200
Volume mm3
16
20
24
Age (Years)

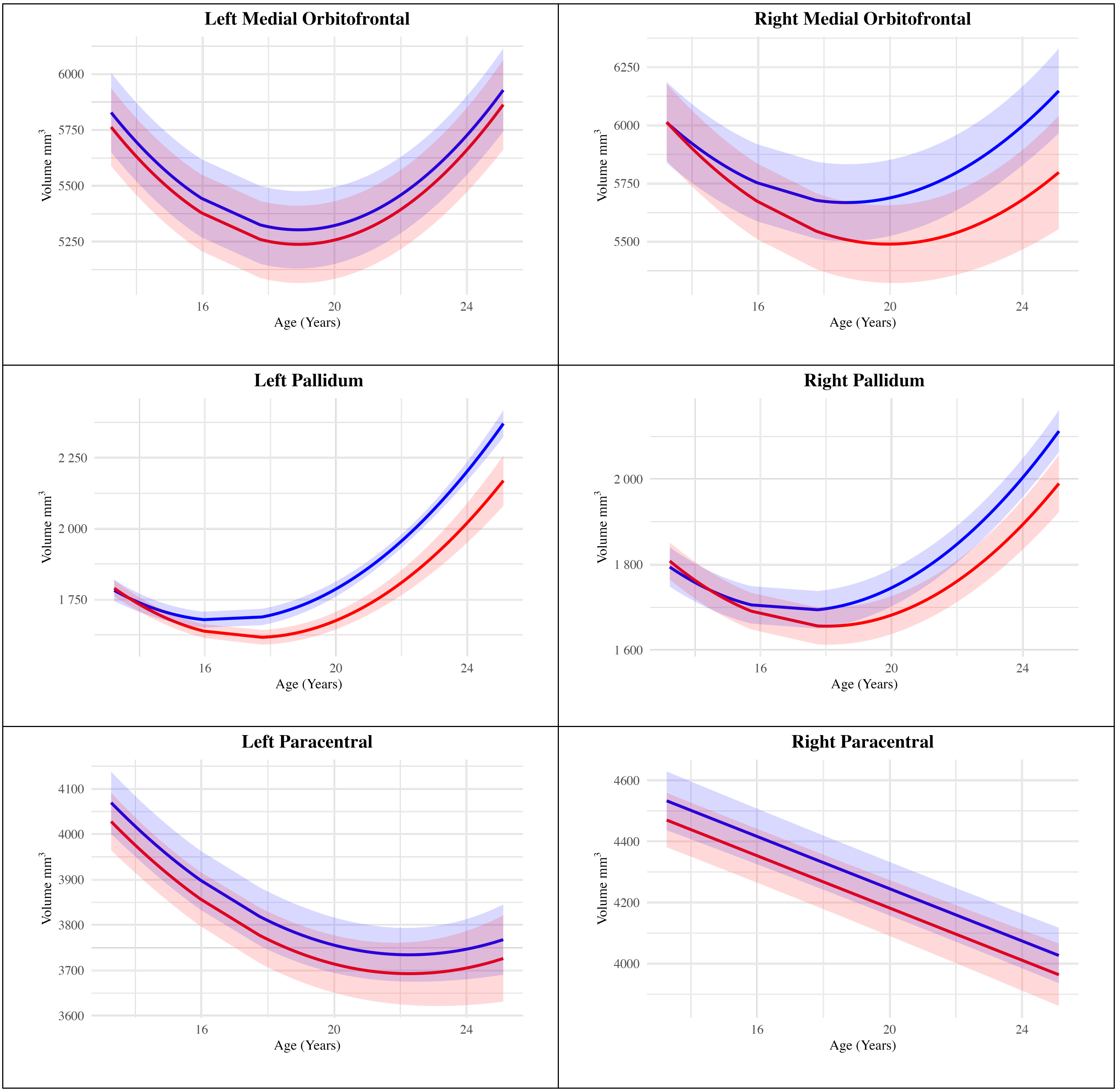
Left Medial Orbitofrontal
6000
5750
5500
5250
Volume mm3
16
20
24
Age (Years)
Right Medial Orbitofrontal
6250
6000
5750
5500
Volume mm3
16
20
24
Age (Years)
Left Pallidum
2 250
2 000
1 750
Volume mm3
16
20
24
Age (Years)
Right Pallidum
2 000
1 800
1 600
Volume mm3
16
20
24
Age (Years)
Left Paracentral
4100
4000
3900
3800
3700
3600
Volume mm3
16
20
24
Age (Years)
Right Paracentral
4600
4400
4200
4000
Volume mm3
16
20
24
Age (Years)

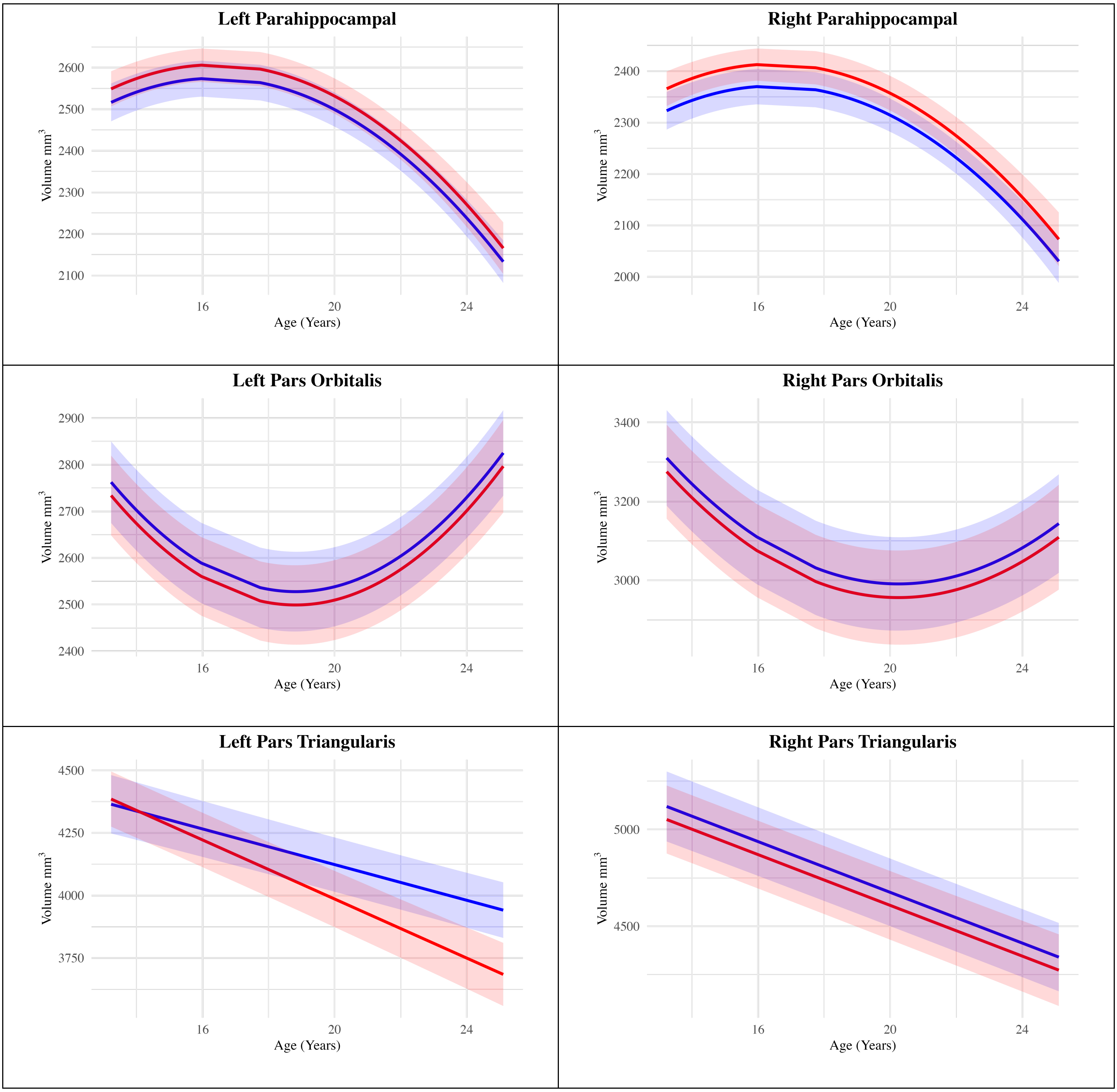
Left Parahippocampal
Right Parahippocampal
Left Pars Orbitalis
Right Pars Orbitalis
Left Pars Triangularis
Right Pars Triangularis
Volume mm3
Age (Years)
16
20
24
2600
2500
2400
2300
2200
2100
2000
2900
2800
2700
3400
3200
3000
4500
4250
4000
3750
5000

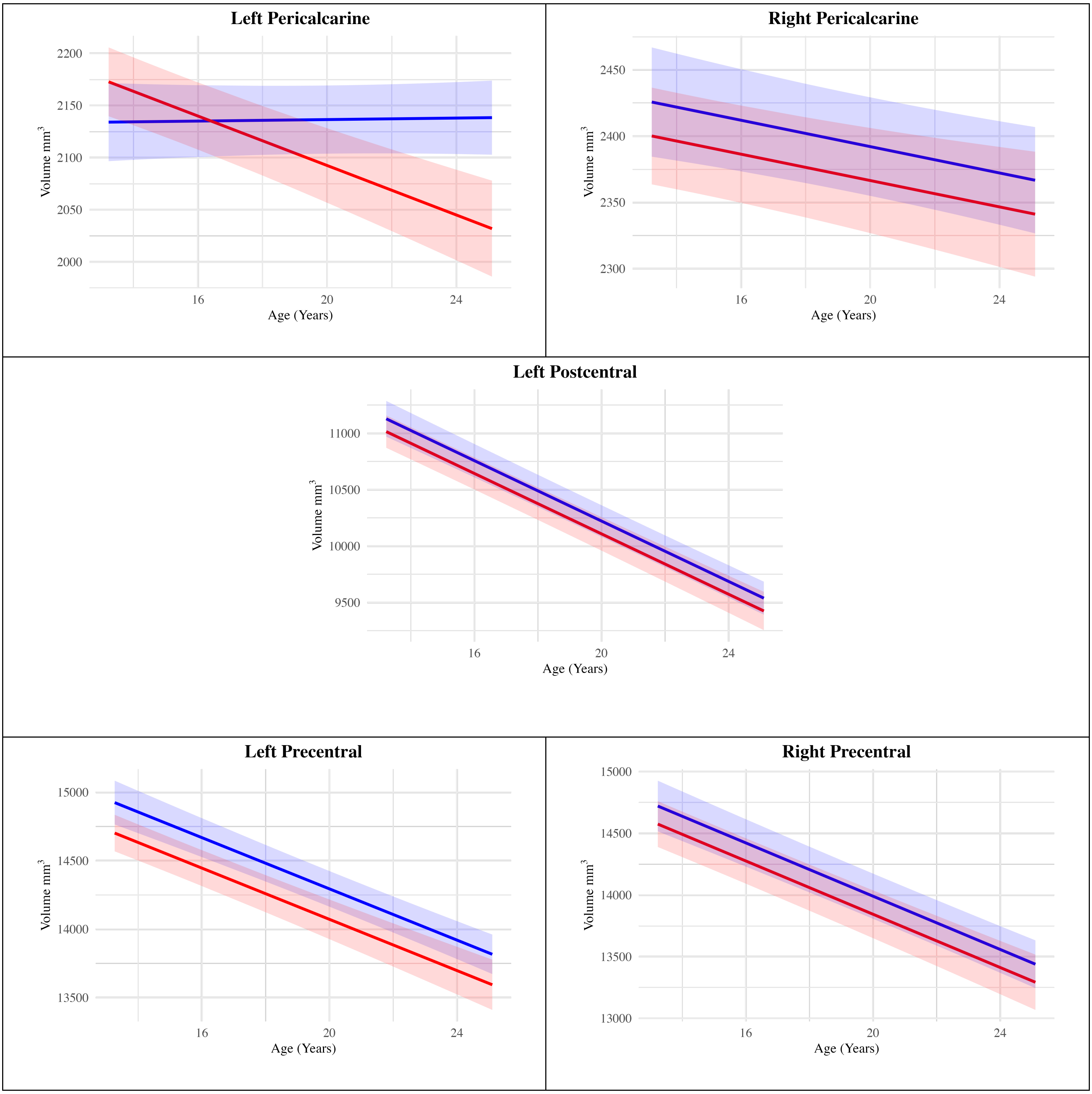
Left Pericalcarine
2200
2150
2100
2050
2000
Volume mm3
16
20
24
Age (Years)
Right Pericalcarine
2450
2400
2350
2300
Volume mm3
16
20
24
Age (Years)
Left Postcentral
11000
10500
10000
9500
Volume mm3
16
20
24
Age (Years)
Left Precentral
15000
14500
14000
13500
Volume mm3
16
20
24
Age (Years)
Right Precentral
15000
14500
14000
13500
13000
Volume mm3
16
20
24
Age (Years)

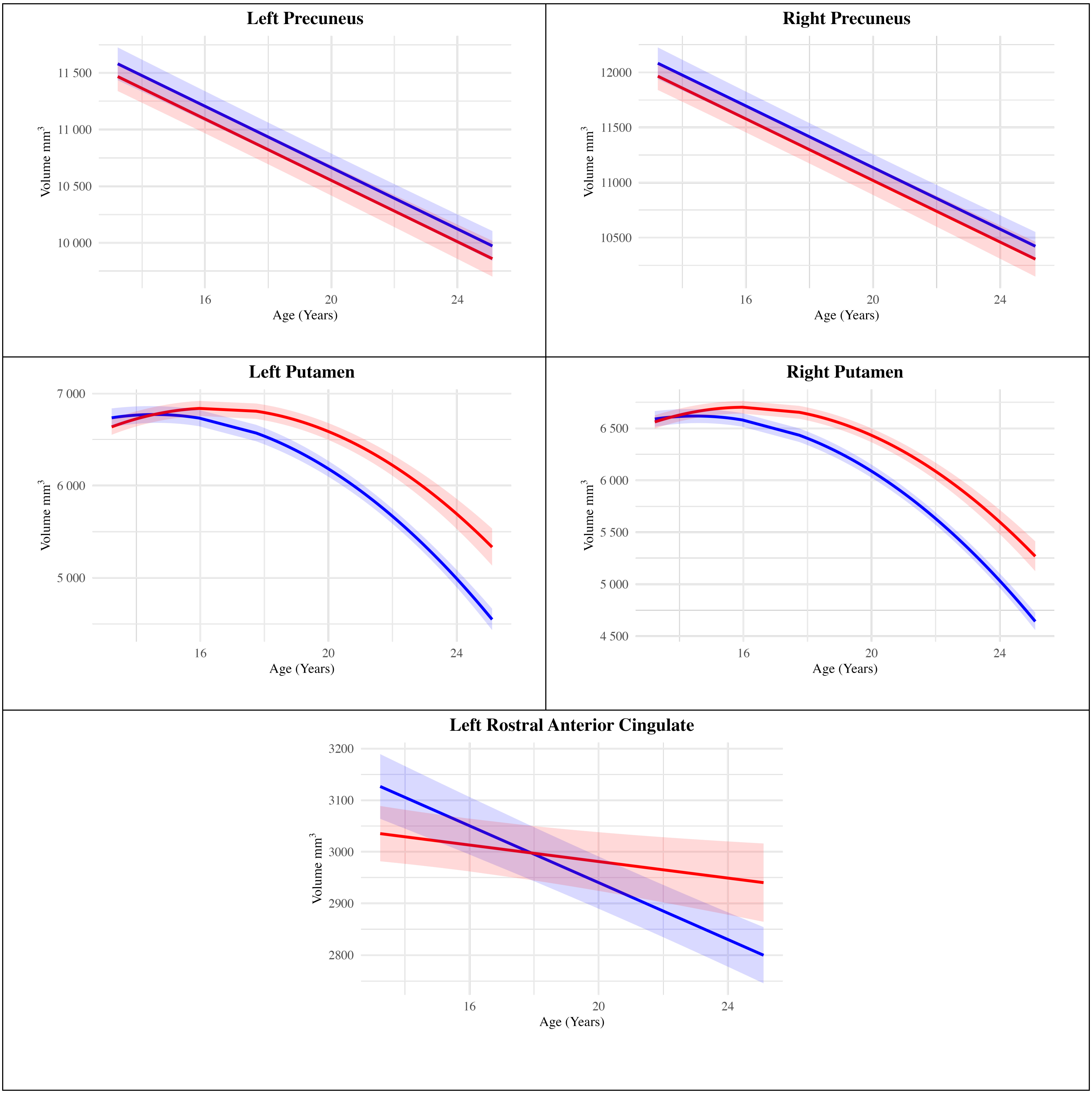
Left Precuneus
11 500
11 000
10 500
10 000
Volume mm3
16
20
24
Age (Years)
Right Precuneus
12000
11500
11000
10500
Volume mm3
16
20
24
Age (Years)
Left Putamen
7 000
6 000
5 000
Volume mm3
16
20
24
Age (Years)
Right Putamen
6 500
6 000
5 500
5 000
4 500
Volume mm3
16
20
24
Age (Years)
Left Rostral Anterior Cingulate
3200
3100
3000
2900
2800
Volume mm3
16
20
24
Age (Years)

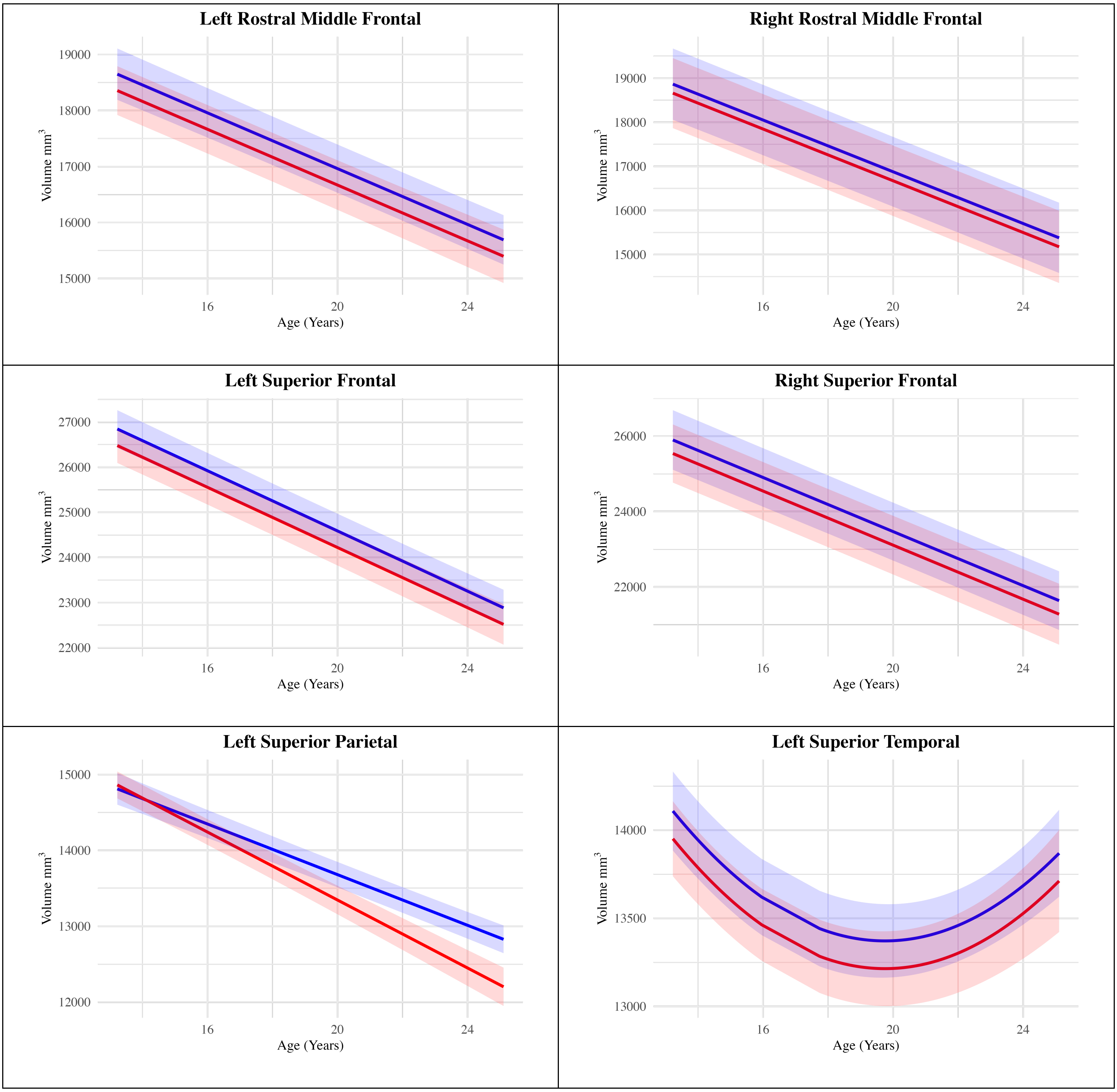
Left Rostral Middle Frontal
19000
18000
17000
16000
15000
Volume mm3
16
20
24
Age (Years)
Right Rostral Middle Frontal
19000
18000
17000
16000
15000
Volume mm3
16
20
24
Age (Years)
Left Superior Frontal
27000
26000
25000
24000
23000
22000
Volume mm3
16
20
24
Age (Years)
Right Superior Frontal
26000
24000
22000
Volume mm3
16
20
24
Age (Years)
Left Superior Parietal
15000
14000
13000
12000
Volume mm3
16
20
24
Age (Years)
Left Superior Temporal
14000
13500
13000
Volume mm3
16
20
24
Age (Years)

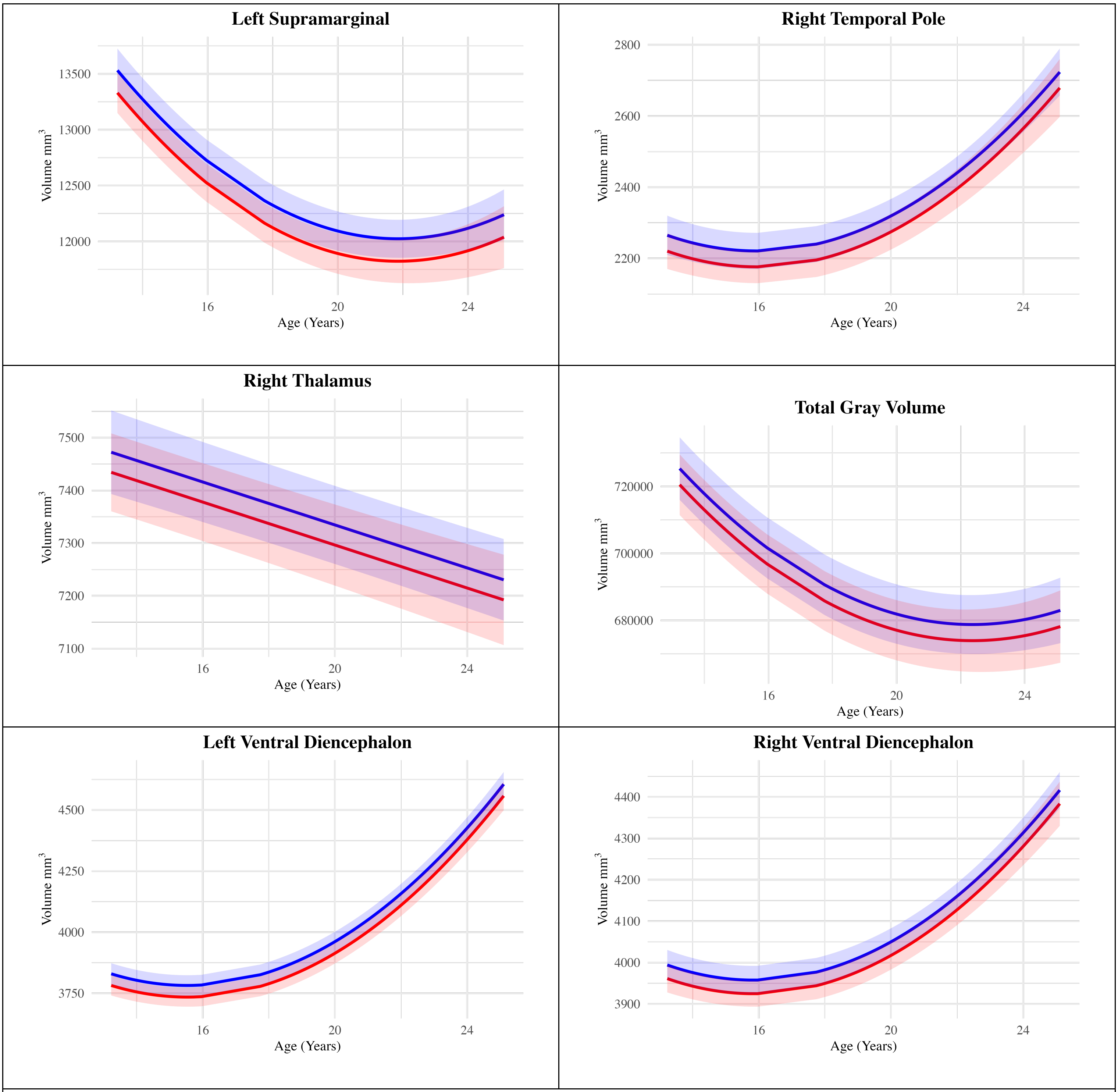


**Legend:** For visualization purposes, two groups were created based on the sum scores from the Olweus Bully/Victim Questionnaire (OB/VQ). The High Bully Victim Group consists of individuals with OB/VQ scores 1 standard deviation above the mean, while the Low Bully Victim Group includes individuals with OB/VQ scores 1 standard deviation below the mean. "Bankssts" refers to the banks of the superior temporal sulcus. The shaded areas represent the 95% confidence intervals for the predicted values.
